# A neurofunctional signature of affective arousal generalizes across valence domains and distinguishes subjective experience from autonomic reactivity

**DOI:** 10.1101/2024.07.17.604003

**Authors:** Ran Zhang, Xianyang Gan, Ting Xu, Fangwen Yu, Lan Wang, Xinwei Song, Guojuan Jiao, Xiqin Liu, Feng Zhou, Benjamin Becker

**Affiliations:** Faculty of Psychology, Southwest University, Chongqing, China; Key Laboratory of Cognition and Personality, Ministry of Education, Chongqing, China; The Center of Psychosomatic Medicine, Sichuan Provincial Center for Mental Health, Sichuan Provincial People’s Hospital, University of Electronic Science and Technology of China, Chengdu, China; School of Life Science and Technology, University of Electronic Science and Technology of China, Chengdu, China; Huaxi MR Research Center (HMRRC), Department of Radiology, West China Hospital of Sichuan University, Chengdu, China; State Key Laboratory of Brain and Cognitive Sciences, The University of Hong Kong, Hong Kong, China; Department of Psychology, The University of Hong Kong, Hong Kong, China

**Author notes:** **Correspondence to** Feng Zhou Southwest University, Tian Sheng RD, No.2, Beibei, ChongQing, 400715, China Mail, Benjamin Becker, The University of Hong Kong Pokfulam, Hong Kong, China Mail.

**Keywords:** Arousal, Valence, Autonomic, Affect, MVPA, Naturalistic paradigm, fMRI

## Abstract

Arousal is fundamental for affective experience and, together with valence, defines the core affective space. Precise brain models of affective arousal are lacking, leading to continuing debates of whether the neural systems generalize across valence domains and are separable from those underlying autonomic arousal or wakefulness. Here, we combined naturalistic fMRI with predictive modeling to develop a brain affective arousal signature (BAAS, discovery-validation design, n=60, 36). We demonstrate its (1) sensitivity and generalizability across mental processes and valence, and (2) neural distinction from autonomic arousal, wakefulness, and stimulation modality (24 studies, n=868). Affective arousal was encoded in distributed cortical-subcortical (e.g., prefrontal, PAG) systems with local similarities in thalamo-amygdala-insula systems between affective and autonomous arousal. We demonstrate application of the BAAS to improve specificity of established valence-specific neuromarkers. Our study provides a biologically plausible model for affective arousal that aligns with the affective space and has a high application potential.

## Introduction

The stream of subjective affective experiences (feelings) represents a continuous assessment of the internal state and its relation with the environment, facilitating adaptive responses, avoiding harm, and identifying opportunities^1^. The mental topology of the highly subjective affective experiences remains controversially debated^2^, but most current theories converge on a dimensional space with core affective domains representing pleasure-displeasure (i.e., valence) and activation-deactivation (i.e., arousal or intensity)^3,4^, which together underlie conscious affective experiences^5,6^. These ‘core affect’ dimensions^5^ represent critical defining features that distinguish affective experiences from other mental states^7^ and form a two-dimensional affective space encompassing specific emotional states (e.g. anger: negative valance and high arousal). Within this core affective space arousal and valance exhibit a V-shaped architecture such that arousal increases in conjunction with either positive or negative valence^8,9^ (see Fig. 1), while increasing levels of intense arousal may progressively overshadow specific emotional states on the subjective, perceptual, and neural level^10,11^.

**Fig. 1.**
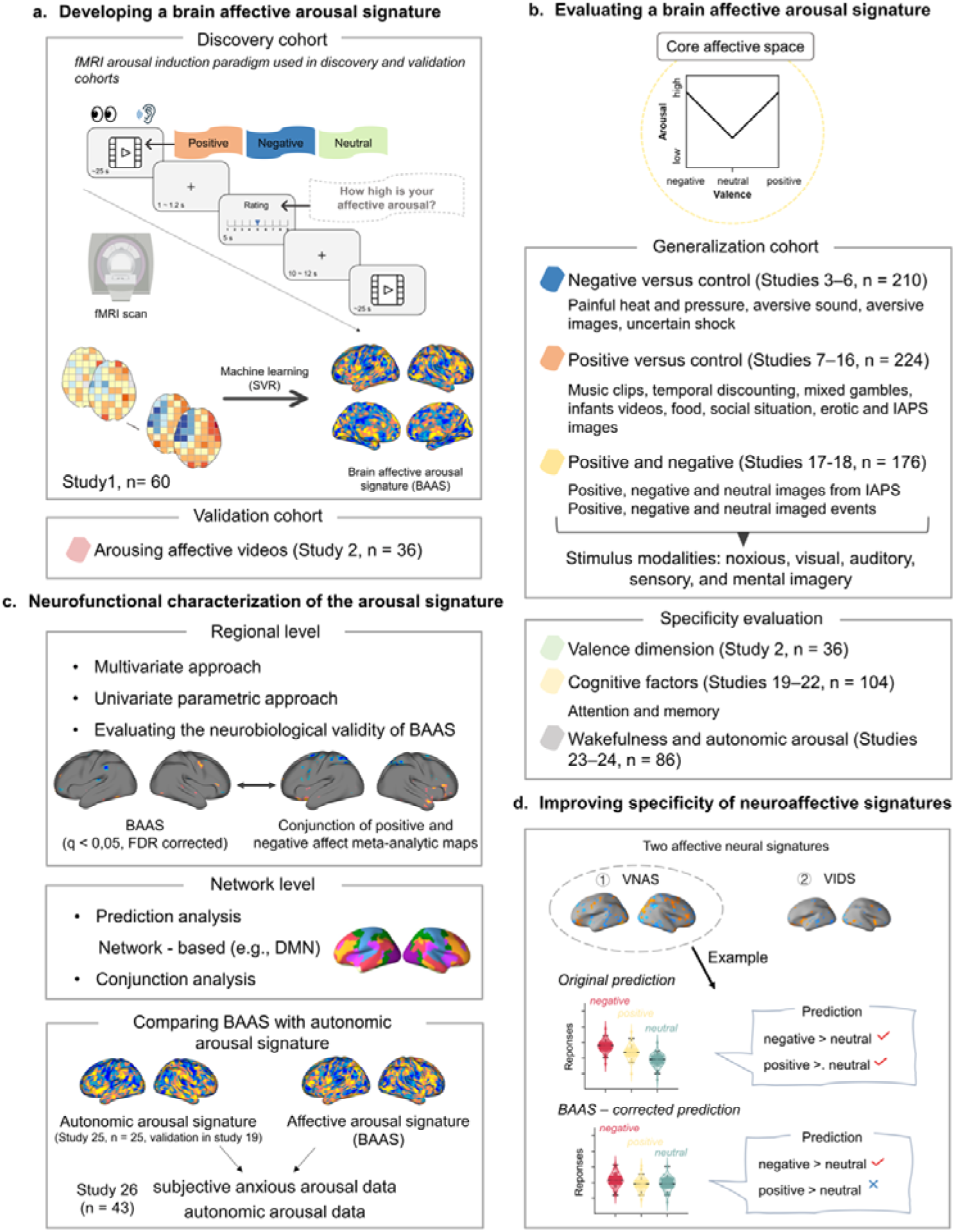
Experimental paradigm and main analyses. (**a**) Building a brain affective arousal signature. A whole-brain model (brain affective arousal signature, BAAS) was developed on the discovery cohort that underwent an arousal induction fMRI paradigm (study 1, n = 60) using the support vector regression algorithm and validated in study 2 (n = 36). (**b**) Evaluating the brain affective arousal signature. Based on the dimensional models of ‘core affect’, BAAS was evaluated the generalizability in sixteen independent datasets (studies 3-18, n = 635) and specificity in seven independent datasets (study 2 and studies 19-22, n = 226). (**c**) Identifying the neurofunctional representation of affective arousal in the brain. Multivariate and univariate approaches were employed to determine the contribution of specific brain systems to predict subjective affective arousal. The thresholded affective arousal signature was compared with the conjunction of meta-analytic maps for positive and negative affect to further validate the biological plausibility of the identified affective arousal brain systems. Prediction analysis and conjunction analysis were next conducted to test performance of isolated brain systems in predicting subjective affective arousal experience. Finally, the neural representations between affective arousal and autonomic arousal signatures were evaluated to validate existed affective arousal-related domains. (**d**) Improving the specificity of neuroaffective signatures. Testing the specificity of two affective neural signatures (VNAS and VIDS) before and after BAAS response corrected. VANS, visually negative affect signature from Čeko et al.^34^; VIDS, visually induced disgust signature from Gan et al.^13^.

The behavioral and neural dynamics of the valance dimension have been extensively mapped at different granularity levels ranging from neural signatures for general negative affect^12^ to signatures that distinguish emotion-specific mental states^13,14^ or subjective affective experiences from accompanying physiological responses^15,16^, respectively. In contrast, and despite its central position in both, classical theoretical accounts and everyday subjective experience of emotion^17,18^ an integrative and generalizable neural model for conscious affective arousal in humans is lacking. The limited conceptualizations and studies to date primarily focus on physiological (autonomic) or wakefulness (vigilance) aspects of arousal^19–21^ rather than the conscious affective experience.

Sophisticated animal studies have established comprehensive and intricate neurobiological models for wakefulness and autonomic arousal, encompassing subcortical systems such as the brainstem, thalamic, and hypothalamic nuclei^22–24^. While there is some overlap between the brain systems controlling wakefulness and hard-wired autonomic reactivity with those involved in affective arousal^23,25,26^, accumulating evidence suggests that the subjective experience that characterizes affective arousal in humans may operate through distinct neural mechanisms^19,27^. In contrast to the anatomical precision of the animal models, human studies on the subjective affective components of arousal have focused on large-scale brain systems (for review see e.g., Satpute et al.^19^), including default mode network subsystems (i.e., anterior medial prefrontal cortex) and the ‘salience network’ (including the anterior cingulate cortex, insula, and amygdala)^28,29^, as well as brainstem regions^30^.

Traditionally, neuroimaging research on affective processes in humans has relied on experimental paradigms employing isolated and sparsely presented stimuli in a single modality (e.g., affective pictures) and analytic approaches that aim at localizing brain regions that show the highest increase in activity by conducting numerous tests across individual ‘voxels’ or regions – strategies characterized by a low ecological validity for dynamic emotional processes in everyday life^31^ and low to moderate effect sizes^32^ and reliability^33^ in terms of characterizing affective mental processes^12,14,16,34^. Recent advances have successfully leveraged neuroimaging combined with machine learning algorithms (i.e., multivariate pattern analysis, MVPA) to develop predictive and more precise and comprehensive neural signatures for subjective affective experiences characterized by a specific emotional state and high negative affective arousal, including general negative affect^12,34^, pain^35^, disgust^13^ , fear^14,15^ and anxious anticipation^16^. The advances allowed to predict the specific subjective affective state with greater effect sizes and robustness compared to traditional local region-based approaches^36,37^ and to demonstrate that the specific emotional states are characterized by common and distinct brain-wide neural representations between subjective emotional states or high affective arousal, respectively^15,16^. This holds the potential to improve the description of mental processes at the brain level^38^. This approach has refined our understanding of affective experiences by demonstrating that they are encoded in distributed brain systems, which is aligned with meta-analytic evidence indicating that affective experiences are linked to activations across a wide array of brain regions^39^.

While initial neuroimaging studies have demonstrated the potential of MVPA to decode stimulus-induced arousal, these studies have not tracked the subjective experience of arousal^40^ nor tested the generalization of the neurofunctional signatures across contexts, socio-cultural diverse samples, states of consciousness or valence, and paradigm classes^41,42^. These aspects form integral components of contemporary affective models, emphasizing the critical significance of subjective experience and appraisal as well as context-dependent affective and neurofunctional computations^43–46^. Very recent conceptualizations further stress the need to segregate neural representations of affective arousal from autonomic arousal and wakefulness, to establish an integrative framework and neural reference space for arousal (e.g., Satpute et al.^19^; for neurofunctional decoding-based support for differentiation in other affective domains see also Taschereau-Dumouchel et al.^15^ and Liu et al.^16^). Despite initial findings demonstrating the promise of neural decoding of arousal, it remains unknown whether affective arousal, a core aspect of affective experiences, exhibits generalizable distributed neural representations in the human brain and whether these representations are independent of valence and distinguishable from the neural representations of physiological arousal and wakefulness.

Moreover, consistent with conceptualizations of the affective space proposing that both positive and negative affect are associated with high levels of arousal, our recent study showed that neural signatures for negative emotions exhibited strong activations in response to positive stimuli compared to neutral ones^47^. This suggests that these signatures may partly capture the arousal inherent to strong affective experiences^48–50^. Together with the observed overshadowing effect of high arousal on the specific emotions across domains^10,11^ this leads to conceptual debates of whether the arousal dimension is represented across valence in the brain and whether accounting for arousal may increase the specificity of decoding specific emotional states from the brain.

Against this background we here aimed to (1) develop a sensitive and generalizable brain signature (i.e., BAAS) that can precisely capture the subjective experience of affective arousal under ecologically valid conditions combining naturalistic functional Magnetic Resonance Imaging (fMRI) with an MVPA-based neural decoding approach, (2) determine whether common neural representations of affective arousal can be determined across the valence domain, multiple senses and modalities, (3) to establish a comprehensive and biologically informed model on how affective arousal is represented in human brain. Given the intricate interaction between autonomic and affective arousal and initial studies demonstrating that the neural representations of autonomic and affective experience might be distinguishable^15,16^ we further examined (4) whether the subjective affective arousal experience is separable from autonomic arousal in humans. Finally, based on previous literature suggesting that high levels of progressively increasing arousal overshadow the representation of specific emotions^11^, we tested if (5) accounting for a precise neural signature of increasing arousal can enhance the specificity of existing affective neural signatures.

To this end, we combined naturalistic fMRI with machine learning-based neural decoding to identify a brain signature predictive of affective arousal intensity in healthy participants (study 1, n = 60) as they experienced a range of positive and negative emotions induced by carefully selected video clips. To enhance the ecological validity of the signature and facilitate immersive and intense arousal experience we utilized naturalistic fMRI experiments using visual-auditory movie clips and requested subjects to report their affective arousal experiences (Fig. 1a). The predictive performance and generalizability of the identified neural signature were assessed in an independent validation cohort (study 2, n = 36), using a modified arousal induction paradigm, with new and longer movie clips covering different aspects of the emotional space. We next employed 16 studies (total n = 610, details see Table S1) to test its generalization and sensitivity across valence domains, modalities, paradigms, populations, and image acquisition systems (Fig. 1b), and additional 6 studies (total n = 190, see Table S1) to evaluate its specificity. The neural representations underlying affective arousal were systematically determined using backward (prediction) and forward (association) encoding models. The neurobiological validity of the BAAS was further examined against meta-analytic maps of negative and positive emotions from large-scale meta-analysis^51^ and the separability from autonomic arousal was determined using a validated decoder for autonomic arousal (fMRI data from Taschereau-Dumouchel et al.^15^, n = 25) (Fig. 1c). Moreover, we investigated whether adjusting for arousal ratings and BAAS responses could enhance the precision of two previously developed and widely employed affective signatures for specific emotional states^13,34^ (Fig. 1d). Overall, this study aims to establish a comprehensive neural model of affective arousal, enhancing our comprehension of how the brain represents affective arousal and refining the prediction specificity of neural signatures for affective experiences.

## Results Behavioral results

In study 1, a total of 40 immersive video stimuli (∼25 s) were employed to induce varying levels of arousal experience. Participants were asked to experience video stimuli immersively and report their current level of affective arousal for each stimulus on a 9-point Likert scale ranging from 1 (very low arousal) to 9 (very high arousal). As shown in Fig. S1a, stimuli induced a wide range of subjective arousal in the discovery cohort which was used to develop the neural signature of subjective affective arousal. Study 2, the validation cohort, employed 28 novel video stimuli 9 (∼60 s), which were entirely different from stimuli used in study 1, to elicit different levels of subjective affective arousal feelings in an independent sample (Fig. S1b).

## Identifying a brain signature for affective arousal

In line with previous studies^12–15^, we employed support vector regression (SVR) in study 1 to develop a brain activity-based signature for subjective affective arousal that predicted the intensity of self-reported arousal experience during the videos. To evaluate the performance of the brain affective arousal signature (BAAS), we conducted the 10 × 10-fold cross-validation procedure in the discovery cohort and additionally validated the signature in the independent study.

The developed BAAS was sensitive to subjective arousal experience in both discovery and independent validation cohorts, as evidenced by significant associations between observed and predicted ratings (Fig. 2a, b). Specifically, in the discovery cohort, the averaged within-subject correlation was 0.88 ± 0.01 (standard error (SE)), the average root mean squared error (RMSE) was 1.36 ± 0.05, the coefficient of determination (R^2^) was [0.61 0.64], the overall (between- and within-subjects) prediction-outcome correlation was 0.82 (R^2^ = 0.65) (Fig. 2a). These findings were replicated in the validation cohort such that BAAS effectively predicted subjective arousal ratings yielding a high prediction-outcome correlation (average within-subject correlation coefficient r = 0.80 ± 0.03 SE, RMSE = 1.68 ± 0.08 SE, R^2^ = 0.42; overall prediction-outcome correlation coefficient r = 0.70, R^2^ = 0.49, Fig. 2b), indicating a sensitive and robust neurofunctional arousal signature. In addition, we categorized video clips in the validation cohort into low (ratings 1, 2, and 3), medium (ratings 4, 5, and 6), and high (ratings 7, 8, and 9) arousal conditions, according to self-reported arousal ratings and applied the two-alternative forced-choice test to examine whether the BAAS could distinguish different arousal conditions. We found that the BAAS response accurately classified high versus moderate (81 ± 6.9%, p = 5.35 ×10^-4^, d = 1.29), moderate versus low (92 ± 4.6%, p = 2.56 ×10^-6^, d = 2.56), and high versus low (100 ± 0.0%, p = 4.66 ×10^-10^, d = 3.21) conditions. Of note, in this analysis, we excluded 4 subjects because they did not have ratings of high or moderate arousal.

**Fig. 2.**
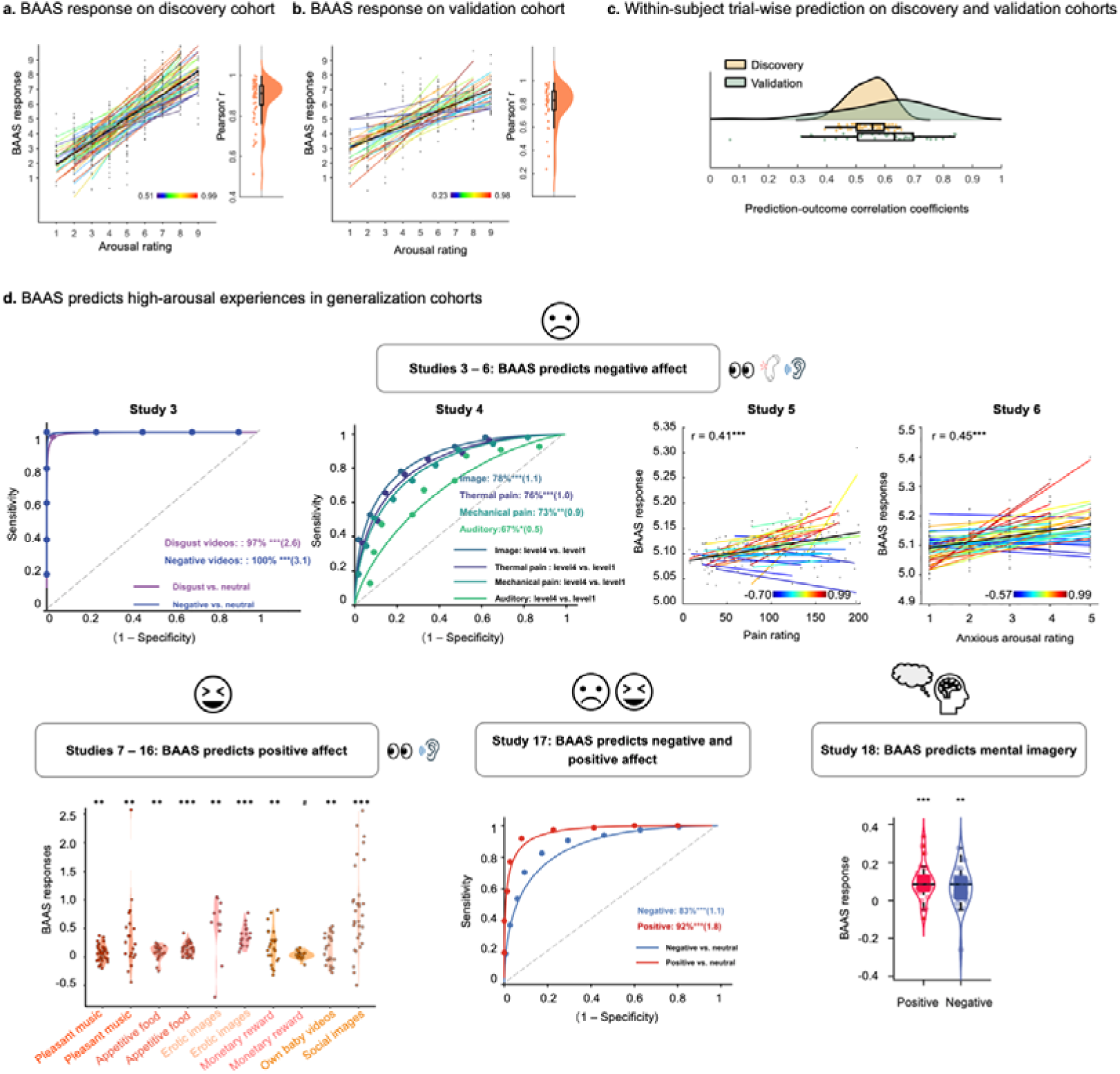
**Evaluating the Brain Affective Arousal Signature (BAAS)**. (**a**) and (**b**) depict predicted arousal experience compared to the actual level of arousal for the cross-validated discovery cohort (study 1, n = 60) and the independent validation cohort (study 2, n = 36), respectively. Colored lines indicate the Pearson correlation coefficients between predicted and true ratings in each subject. The black line reflects the overall correlation. Raincloud plots show the distribution of correlation. (**c**) Distribution of within-subject trial-wise prediction-outcome correlation coefficients for each participant in the discovery and validation cohorts, respectively. Yellow represents correlation coefficients in the discovery cohort, while green represents correlation coefficients in the validation cohort. (**d**) BAAS accurately predicts (forced-choice classification) or strongly responds (one-sample t test) to high arousal experiences induced by stimuli across multiple modalities. Upper panel: BAAS accurately predicts high arousing negative affect in studies 3 – 6 (total n = 210, prediction performance is shown as forced-choice classification accuracy (Cohen’s d) or prediction-outcome correlation); Left lower panel: BAAS (marginally) significantly responds to high arousing positive affect in studies 7 – 16 (total n = 224); Middle lower panel: BAAS accurately predicts high arousing negative and positive affect in study 17 (total n = 150, prediction performance is shown as forced-choice classification accuracy (Cohen’s d)). Right lower panel: BAAS accurately predicts high arousing positive and negative imaged events in study 18 (total n = 26). See Table S1 for the details of each contrast. # Marginally significant, * P < 0.05, ** P < 0.01, *** P < 0.001. BAAS, Brain affective arousal signature.

Given that previous studies showed activity in the visual cortex in response to arousing emotional scenes, suggesting that visual processing contributes to arousal^52–55^, we tested to which extent the BAAS depended on the visual cortex. Therefore, we retrained the decoder excluding the entire occipital lobe. We observed similar performance including significant prediction-outcome correlations and coefficient of determination in this modified setup, suggesting that the prediction is not solely reliant on emotional schemas encoding within visual regions (further details see Supplementary Results and Fig. S2).

## Within-subject trial-wise prediction

Affective arousal experience is a highly subjective state and can rapidly fluctuate within a short period, typically on the order of minutes or even seconds^27,56,57^. We next examined to what extent the population-level BAAS can accurately predict individual-level variations in arousal, specifically on a trial-by-trial basis. To this end, the single-trial analysis was used to obtain individual-specific trial-wise activation maps. Subsequently, the BAAS pattern expressions of these single-trial activation maps were calculated and then correlated with corresponding true ratings for each subject separately. In the discovery cohort, the BAAS accurately predicted individual arousal experience (n = 40 trials; within-participant r = 0.55 ± 0.01 SE, 10×10 cross-validated; Fig. 2c). Of note, the BAAS successfully predicts trial-by-trial arousal ratings in all participants within the discovery cohort (all p ≤ 3.8×10^-3^). We additionally validated the predictive performance of the BAAS (developed on data from the discovery cohort) on the validation cohort with respect to predicting single trial responses in each participant in the discovery cohort. The predictive performance of the BAAS on the validation cohort was similar to those observed in the discovery cohort (n = 28 trials; within-participant r = 0.60 ± 0.03 SE, Fig. 2c), and the BAAS significantly predicted trial-wise affective experience for over 88.8% participants in the validation dataset. Taken together, our findings suggest that BAAS accurately predicts the level of momentary arousal feelings on the individual level.

## Evaluating generalization of BAAS in independent datasets

We examined whether the BAAS could capture affective arousal across valence, various stimulus modalities and different experimental approaches. A substantial body of evidence suggests a V-shaped relationship between affective valence and arousal levels, wherein individuals experiencing more intense positive or negative emotions generally report heightened arousal^9^ (Fig. 1). We first determined whether the BAAS distinguishes high arousal negative experiences from low arousal negative experiences across affective domains, stimulus modalities and generalizes to the stimulus-free state (aversive anticipation) (Fig. 2d). Study 3 employed a naturalistic affective video-watching fMRI paradigm involving strong disgust-inducing, arousal-matched negative, and neutral video stimuli (affective ratings see Table S2). We found that the BAAS successfully classified disgust and negative from neutral experiences with high accuracies (disgust versus neutral: accuracy = 97 ± 1.8% SE, p < 1.00 ×10^-10^, d = 2.64, negative versus neutral: accuracy = 100 ± 0.0% SE, p < 1.00 ×10^-10^, d = 3.08) supporting its generalizability (Fig. 2d ‘Study 3’). Of note, the BAAS could not classify arousal matched negative (arousal: mean ± SD = 7.15 ± 0.41) from disgust experience-specific neural activity in this dataset (arousal: mean ± SD = 7.23 ± 0.50) (accuracy = 60 ± 5.5% SE, p = 0.09, d = 0.31, Fig. S3), further supporting that the signature captures arousal across negative affective states. Study 4 induced negative experience with four types of aversive stimuli—painful heat, painful pressure, aversive images and aversive sounds^34^. The BAAS was able to effectively distinguish between high and low levels (i.e., level 4 and level 1) of aversive image (accuracy = 78 ± 5.6% SE, p = 3.31 ×10^-5^, d = 1.08), thermal pain (accuracy = 76 ± 5.7% SE, p = 1.14 ×10^-4^, d = 0.96), mechanical pain (accuracy = 73 ± 6.0% SE, p = 1.00 ×10^-3^, d = 0.90), and aversive sound (accuracy = 67 ± 6.3% SE, p = 0.01, d = 0.48) (Fig. 2d ‘Study 4’). Additionally, the BAAS could accurately predict the aversive ratings induced by aversive image (r = 0.30, p = 6.70 ×10^-6^), thermal pain (r = 0.24, p = 4.09 ×10^-4^), mechanical pain (r = 0.40, p = 6.04 ×10^-10^), and aversive sound (r = 0.16, p = 0.02). These results indicated that the BASS generalizes to pain-associated states and across stimulus modalities. In line with the extension to pain assessment, in study 5^58^, the BAAS successfully predicted subjective pain ratings (prediction-outcome r = 0.41, p = 2.15 ×10^-9^) as well as accurately discriminated high levels of pain versus low levels of pain (44.8°C vs. 48.8°C) (accuracy = 85 ± 6.2% SE, p = 6.62 ×10^-5^, d = 1.16) (Fig. 2d ‘Study 5’). Moreover, we further tested whether the BAAS is independent of the actual stimulus exposure by testing generalization to data from an ‘Uncertainty-Variation Threat Anticipation’ (UVTA) task in study 6 ^16^. During this task participants anticipate aversive electrical stimulation with varying levels of uncertainty and retrospectively rated their anxious arousal during the anticipation period. The BAAS accurately predicted the subjective anxious experience with prediction-outcome r = 0.45 (p < 1.00 ×10^-10^) (Fig. 2d ‘Study 6’), confirming its generalization to stimulus-free internal mental processes. Together, these findings demonstrate that within the negative affect domain, the BAAS shows strong generalizability across stimuli and modalities, and this generalizability also extends to affective experiences that do not involve animate stimuli, such as noxious heat and pressure.

We next determined generalizability of the BAAS in the positive domain by capitalizing on data from 10 independent studies, employing different strategies and modalities to induce high arousing positive affect across the experience and anticipation period (e.g., music, images of appetizing food, erotic images, cues of monetary rewards, and socially relevant stimuli with 2 studies in each category)^41,42,59–66^. The results indicated that BAAS exhibited significantly stronger responses to positive compared to control conditions in nine out of ten studies (refer to Table 1 and Fig. 2d ‘Studies 7-16’), and a marginally significant stronger response (p = 0.07) in the remaining study, suggesting that the BAAS can capture high-arousal positive experiences across stimulus modalities (i.e., vision and auditory). Importantly, in line with the findings in the negative affect domain, the BAAS showed well predictive performances in positive affective experiences induced by both animate stimuli (e.g., erotic images) and inanimate objects (e.g., cues of monetary rewards).

**Table 1.**
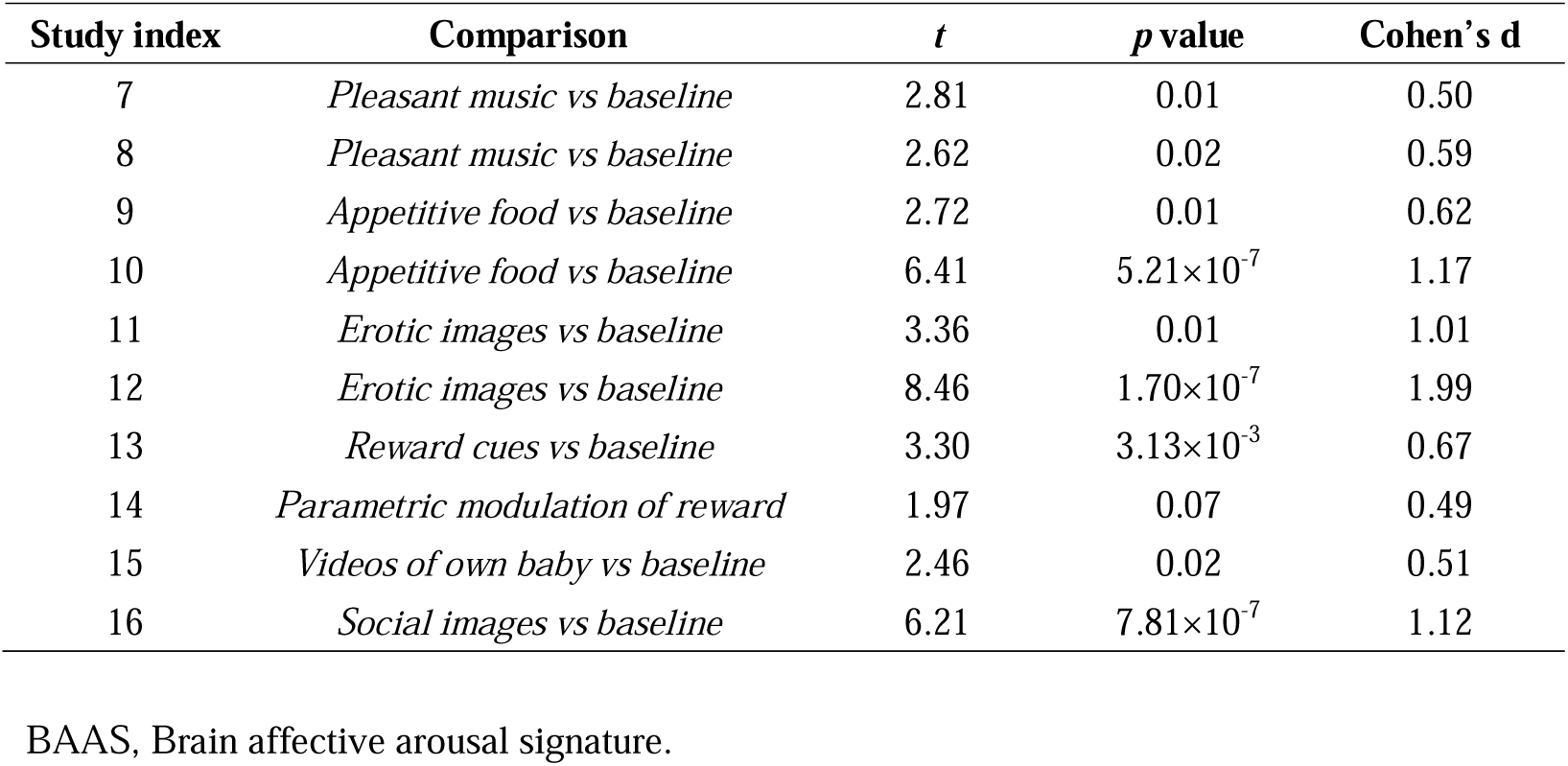
Generalization performance of the BAAS on studies 7– 16.

| Study index | Comparison | *t* | *p* value | Cohen's *d* |
| --- | --- | --- | --- | --- |
| 7 | *Pleasant music vs baseline* | 2.81 | 0.01 | 0.50 |
| 8 | *Pleasant music vs baseline* | 2.62 | 0.02 | 0.59 |
| 9 | *Appetitive food vs baseline* | 2.72 | 0.01 | 0.62 |
| 10 | *Appetitive food vs baseline* | 6.41 | $5.21 \times 10^{-7}$ | 1.17 |
| 11 | *Erotic images vs baseline* | 3.36 | 0.01 | 1.01 |
| 12 | *Erotic images vs baseline* | 8.46 | $1.70 \times 10^{-7}$ | 1.99 |
| 13 | *Reward cues vs baseline* | 3.30 | $3.13 \times 10^{-3}$ | 0.67 |
| 14 | *Parametric modulation of reward* | 1.97 | 0.07 | 0.49 |
| 15 | *Videos of own baby vs baseline* | 2.46 | 0.02 | 0.51 |
| 16 | *Social images vs baseline* | 6.21 | $7.81 \times 10^{-7}$ | 1.12 |

BAAS, Brain affective arousal signature.

These findings were further replicated in a large dataset with arousal-matched negative and positive affect induction^67^ (n = 150, Fig. 2d ‘Study 17’), where the BAAS accurately classified negative and positive from neutral conditions (negative vs. neutral: accuracy = 83 ± 3.1% SE, p < 1.00 ×10^-10^, d = 1.12; positive vs. neutral: accuracy = 92 ± 2.2% SE, p < 1.00 ×10^-10^, d = 1.78). Overall, BAAS showed remarkable generalizability across valence domains and multiple modalities.

Finally, we evaluated BAAS response in an imagination task (Fig. 2d ‘Study 18’). The task required participants to imagine various scenarios, including both positive and negative events during fMRI scanning^68^. We compared the neural responses to these positive and negative imagined events to a baseline condition. The BAAS exhibited significantly stronger responses to both positive (t = 4.97, p = 4.03×10^-5^, d = 0.97) and negative (t = 3.32, p = 2.80×10^-3^, d = 0.65) imagined events relative to the baseline. These results demonstrate that the BAAS, originally identified based on responses to external stimuli, is also capable of predicting internally generated affective experiences.

Notably, these sixteen datasets were collected with diverse MRI scanners, populations and were preprocessed using various pipelines. Our findings collectively highlight the robust generalizability and high sensitivity of the BAAS originally developed in the movie-watching paradigm in effectively predicting high affective arousal experiences across valence domains, diverse stimuli, modalities, MRI systems, preprocessing pipelines, and populations. Moreover, the BAAS is not limited to capturing animate objects in the affective domain (more evidence see Supplementary Results).

## Validating specificity of BAAS in independent datasets

To sufficiently consider the specificity of BAAS, we conducted tests on BAAS in a series of independent datasets. We first tested whether valence, another core dimension of affective experience, contributes to BAAS predictions. Given that both positive and negative video clips induced high arousal in the validation cohort (positive: mean ± SD = 7.37 ± 0.35; negative: mean ± SD = 7.94 ± 0.29, details of videos see Table S2), we hypothesized that if BAAS primarily captures arousal rather than valence information, it should accurately classify high arousing positive and negative video clips from low arousing neutral video clips, yet it might not differentiate between the negative and positive video clips. Consistent with our hypotheses, BAAS demonstrated significant discriminability of unspecific high arousal experiences (positive versus neutral: 97 ± 2.7%, p = 1.08×10^-9^, d = 2.63; negative versus neutral: 97 ± 2.7%, p = 1.08×10^-9^, d = 2.90) and poor accuracy in discriminating between two high-arousal experiences (negative versus positive: 53 ± 8.3%, p = 0.87, d = 0.04) (Fig. S4), suggesting high specificity of the BAAS for capturing arousal across valence.

Next, to estimate the effect of potential confounding factors (e.g., attention, or working memory), we examined BAAS performances during several effortful cognitive tasks (studies 19 – 22). Specifically, we assessed the BAAS response to incongruent and congruent trials with low levels of discriminability in both the Eriksen Flanker task^69^ (t = -0.16, p = 0.87, d = 0.07) and the Simon task^70^ (t = 0.34, p = 0.74, d = -0.04) (Fig. S5a). Additionally, the BAAS did not differentiate 1-back blocks from the baseline in the N-back task^71^ (accuracy = 67 ± 8.6% SE, p = 0.10, d = 0.72) (Fig. S5b). During the emotional memory task^72^, the BAAS successfully classified remembered positive and negative items from remembered neutral items (remembered positive versus remembered neutral: 89 ± 6.0%, p = 4.92×10^-5^, d = 2.24; remembered negative versus remembered neutral: 96 ± 3.6%, p = 4.17×10^-7^, d = 2.17) (Fig. S5b), suggesting that affective arousal decoder is not reliant on memory processes but specifically captures arousal across different emotional valences.

To further verify that this affective arousal signature is dissociable from wakefulness and autonomic arousal, we examined its discriminability during a sleep-wake study with concomitant fMRI- electroencephalogram (EEG) (study 23) and Pavlovian threat conditioning task (study 24). In the sleep-wake study^73^, 33 participants had two 10-minute resting-state sessions and several sleep sessions, with simultaneously collected fMRI and EEG, and exhibited both sleep and wake stages (details see Supplementary Methods). Of these, 28 participants with complete data were included, and the BAAS was applied to their time series to obtain mean pattern responses for sleep and wake stages. Results showed that the BAAS could not significantly differentiate wakefulness from sleep (accuracy = 54 ± 9.4% SE, p = 0.85, d = -0.27) (Fig. S6a). In the Pavlovian threat conditioning task^74^, simultaneously acquired fMRI and psychophysiological arousal responses (i.e., galvanic skin response, GSR) were recorded for each participant. Consistent with our previous study^74^, we only included participants (n = 58) who exhibited stronger psychophysiological threat responses (i.e., mean CS+ > mean CS-, conditioned stimulus, CS) during threat acquisition. The BAAS was unable to discriminate CS+ (associated with high GSR) from CS- (associated with low GSR) (accuracy = 60 ± 6.4% SE, p = 1.15, d = 0.49, see Fig. S6b).

Together, these findings consistently suggest that the identified BAAS is specifically sensitive to affective arousal and dissociable from valence, cognitive demand, wakefulness, and autonomic arousal.

## The neurofunctional characterization of the affective arousal signature

To understand the neurofunctional core system of affective arousal in humans we carefully determined the brain regions reliably contributing to the BAAS prediction performance. To this end, we conducted bootstrap tests to identify voxels that most reliably contribute to the prediction (q < 0.05, false discovery rate (FDR) corrected). As shown in Fig. 3a, activity in amygdala, periaqueductal gray (PAG), parahippocampal gyrus, lateral orbitofrontal cortex (lOFC), dorsomedial prefrontal cortex (dmPFC), subgenual anterior cingulate (sACC), and superior frontal gyrus (SFG) strongly predict increased affective arousal. Conversely, most reliable negative weights (i.e., higher arousal with decrease activity) were found in the posterior insula / rolandic operculum, supplementary motor area (SMA), middle frontal gyrus (MFG), inferior parietal lobule (IPL) and posterior cingulate cortex (PCC). These regions have been reported in the literature for a range of highly arousing emotional experiences. Moreover, parcellation-based analysis was employed to determine local brain regions that were predictive of affective arousal. Specifically, for each of the 275 brain regions^75,76^, we trained and cross-validated parcel-wise models predictive of affective arousal in the discovery dataset and further tested them in the independent validation dataset. As shown in Fig. 3b, affective arousal experience could be significantly predicted by activations in widely distributed regions (p < 0.001, uncorrected), which overlapped with a number of areas in BAAS patterns thresholded at q < 0.05 (FDR corrected, 5000-sample bootstrap).

**Fig. 3.**
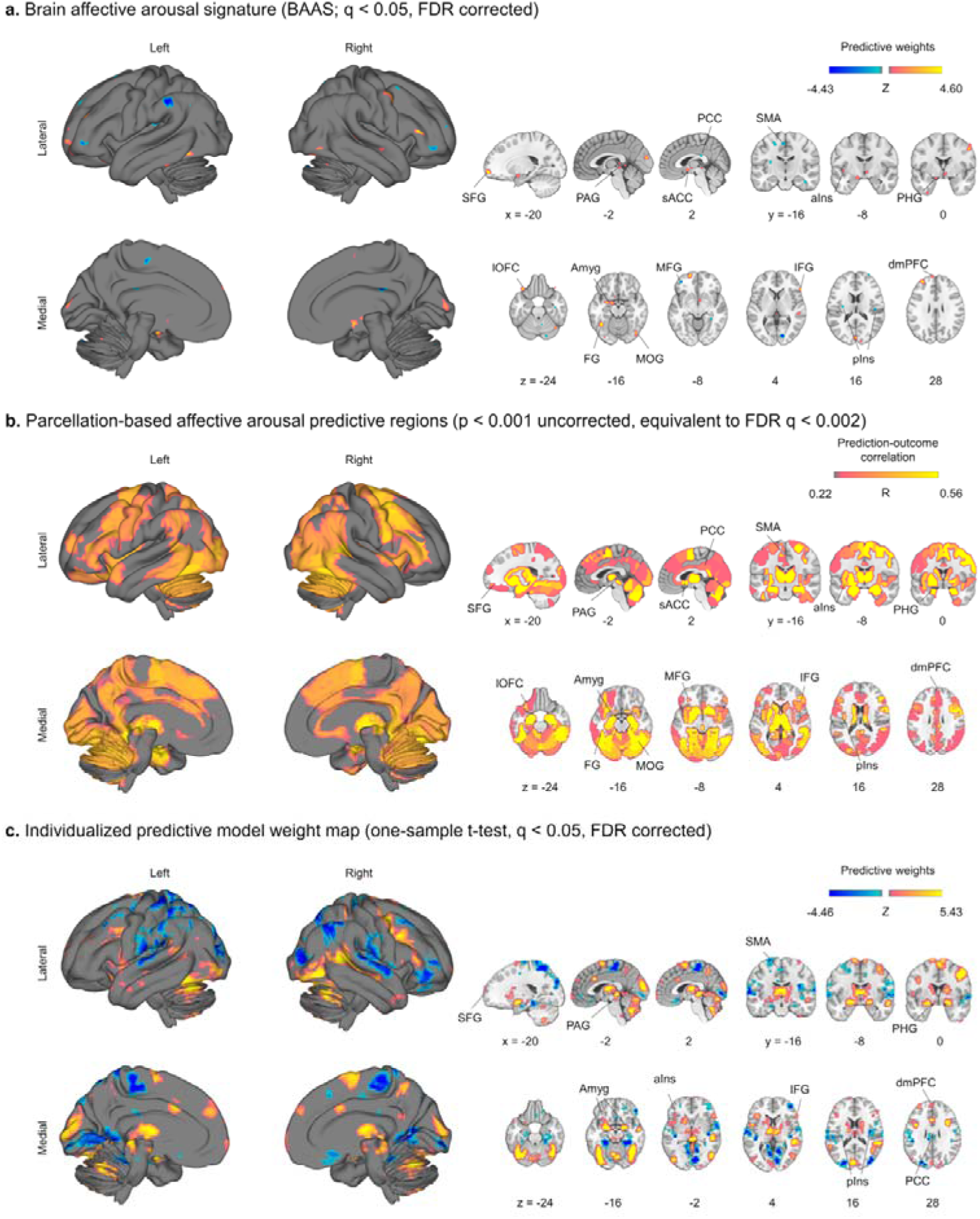
Brain systems for affective arousal. (**a**) shows the thresholded BAAS weight maps based on 5,000 samples bootstrap test (q < 0.05, FDR corrected). (**b**) displays brain regions that significantly (p < 0.001 uncorrected, equivalent to FDR q < 0.002) predict affective arousal experience in both discovery (10×10-fold cross validation) and validation cohorts revealed by parcellation-based analysis. (**c**) summarizes multivariate patterns trained on individual subjects and depicts brain regions that consistently predict subjective affective arousal across n = 60 participants (the discovery cohort) using a one-sample t test (q < 0.05, FDR corrected) based on training a separate model for each subject’s trial by trial ratings. In (**b**), hot color indicates the averaged prediction-outcome correlation values from the discovery cohort and validation cohort. In (**a**) and (**c**), hot color indicates positive values, whereas cold color indicates negative values. BAAS, Brain affective arousal signature; aIns, anterior insula; Amyg, amygdala; dmPFC, dorsomedial prefrontal cortex; FG, fusiform gyrus; IFG, inferior frontal gyrus; lOFC, lateral orbitofrontal cortex; MFG, middle frontal gyrus; MOG, middle occipital gyrus; PAG, periaqueductal gray; PCC, posterior cingulate cortex; PHG, parahippocampal gyrus; pIns, posterior insula; sACC, subgenual anterior cingulate; SFG, superior frontal gyrus; SMA, supplementary motor area^38^.

Given that affective experience is a highly subjective and individually constructed state, with significant variations among individuals^77^, we further developed a predictive model for each participant based on single-trial data and performed a one-sample t-test analysis (treating participants as a random effect) on the weights derived from individualized multivariate predictive models (q < 0.05, FDR corrected). This approach is based on separate linear SVR prediction analysis for each participant in the discovery cohort using their single-trial activation maps and the one-sample t-test allows to identify arousal-predictive regions consistently engaged across participants. In line with the population-level BAAS model, we observed that amygdala, parahippocampal gyrus, PAG, anterior insula, dmPFC, and SFG robustly predict enhanced affective arousal, while reductions in affective arousal were strongly linked to signals from posterior insula, SMA, MFG, PCC (Fig. 3c). Notably, brain regions associated with affective arousal determined with univariate parametric modulation approach identified a similar set of broadly distributed regions (see Supplementary Results and Fig. S7).

To further validate the identified biologically plausible affective arousal brain system, we additionally compared the thresholded BAAS (q < 0.05, FDR corrected) with the conjunction of the meta-analytic maps of ‘negative’ and ‘positive’ affective experiences which were obtained from the Neurosynth (association test q < 0.05, FDR corrected, maps see Fig. S8). Given that feeling of arousal are more likely to be accompanied by valanced feelings, positive or negative^9^, we hypothesized that the overlapping between these two meta-analytic maps might share common brain regions with the BAAS. Consistent with our prediction, several overlapping regions (e.g., amygdala, parahippocampal gyrus, including inferior frontal gyrus (IFG), lOFC, and sACC) between two patterns were observed using conjunction analyses (Fig. 4a). As shown in Fig. S8, brain regions including IFG, SFG, dmPFC, amygdala, parahippocampal gyrus, and insula positively associated with both ‘negative’ and ‘positive’ meta-analytic maps were also found significant in the BAAS, though there were not directly overlapping. Conversely, SMA and precentral gyrus regions were negatively associated with both ‘negative’ and ‘positive’ meta-analytic maps and were reliably predictive of decreased arousal in the BAAS. These findings offer evidence confirming the identified regions involved in affective arousal processing.

**Fig. 4.**
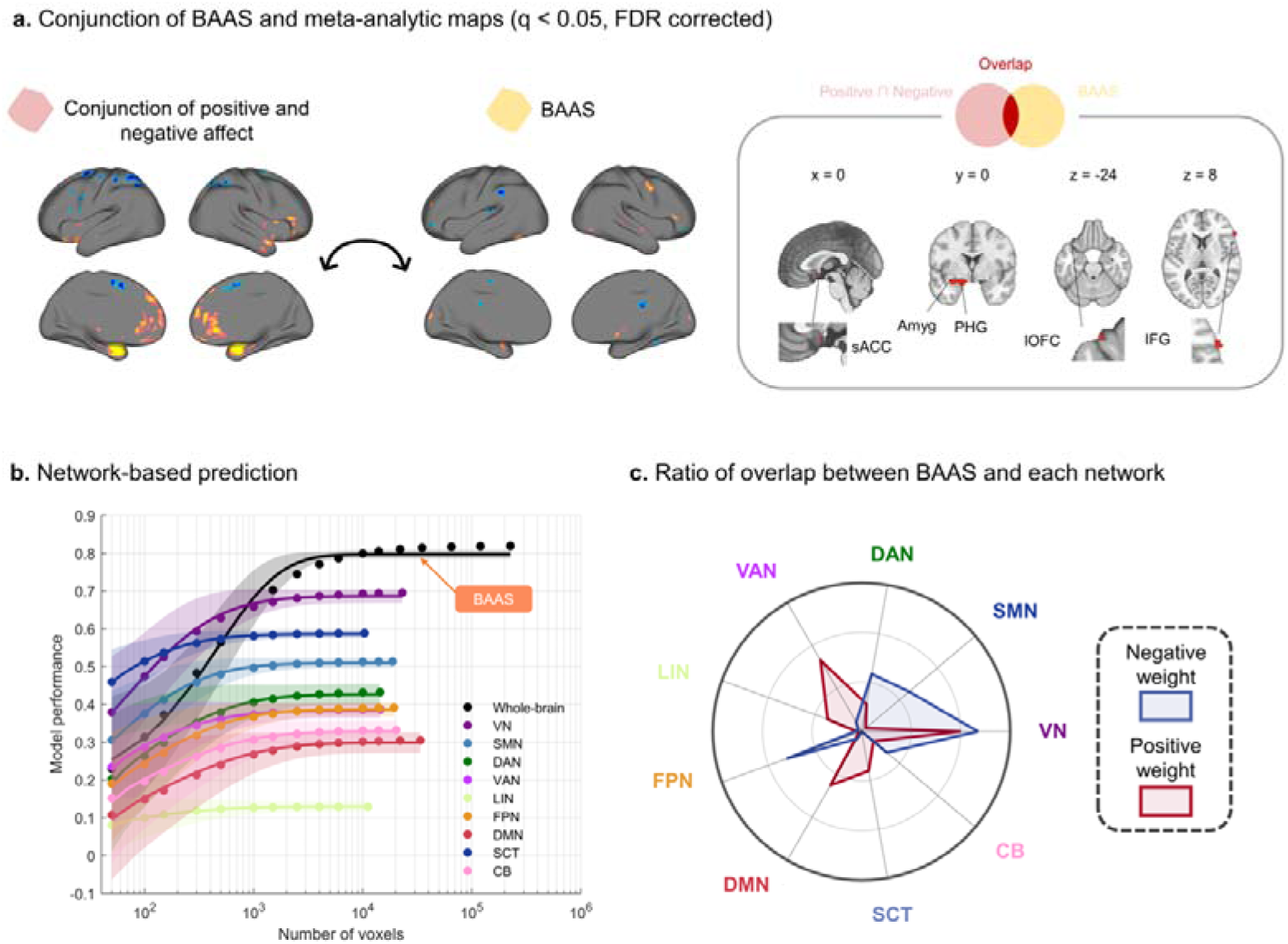
The distributed neural representations of affective arousal exhibited neurobiological plausibility. (**a**) displays overlapping areas between thresholded BAAS and the conjunction of the meta-analytic maps of ‘negative’ and ‘positive’ affect from Neurosynth. (**b**) shows that the performance of the model was evaluated by using an increasing number of voxels/features (on the x-axis) to predict subjective affective arousal in different regions of interest, such as the whole brain (in black), cerebellar brain (in pink), subcortical regions (in dark blue) or large-scale resting-state networks. The y-axis depicts the cross-validated correlation between predicted and actual outcomes. The colored dots demonstrate the average correlation coefficients, while the solid lines indicate the average parametric fit, and the shaded regions reflect the standard deviation. The model’s performance is optimized by randomly sampling approximate 10,000 voxels across the entire brain. (**c**) shows the relative proportions of the voxels of thresholded BAAS within each large-scale functional network given the total number of voxels within each network. Red represents voxels with positive weight in the BAAS, while blue represents voxels with negative weight. BAAS, Brain affective arousal signature; VN, Visual network; SMN, Somatomotor network; DAN, Dorsal attention network; VAN, Ventral attention network; LIN, Limbic network; FPN, Frontoparietal network; DMN, Default mode network; SCT, Subcortical network; CB, Cerebellar brain.

Together, our results identify the conscious experience of affective arousal is represented in distributed subcortical and cortical regions involved in salience information processing (e.g., amygdala, insula)^78,79^, emotional awareness (e.g., frontal regions)^80^, and emotional memory (e.g., hippocampus)^81^, as well as areas that have been emphasized as a core system mediating the arousal function in animal models (brainstem)^19^.

## Alternative models to determine the contribution of isolated arousal-prediction models: BAAS predicts more precise that isolated networks

Previous studies have emphasized contributions of DMN, salience and subcortical networks to affective arousal processing^28,29^ and psychological construction theories of emotion propose arousal as one component of ‘core affect’ underlying functions that are neurobiologically supported by large-scale networks^39^. To address this question, we obtained the predictive SVR model by restricting it to masks corresponding to (1) each of seven large-scale resting-state functional networks, (2) a subcortical network, and (3) the cerebellum (see Methods session) in the discovery cohort. This allowed us to examine the extent to which these models could predict arousal experiences compared to the whole-brain BAAS. The results showed that brain networks, especially the visual network and subcortical network, to some extent, predict subjective affective arousal ratings (see Fig. 4b and Table S3). Considering the potential effects of the number of features/voxels in prediction analyses, as shown in Fig. 4b, we implemented a control procedure. Specifically, we conducted 5,000 repeated random voxel selections from different brain regions, including the entire brain, subcortical, or individual resting-state networks (averaged over 5,000 iterations). This approach was carried out to assess the impact of varying the sampled brain systems. Interestingly, our results revealed that the asymptotic prediction achieved when sampling from all brain systems, as demonstrated in the BAAS (black line in Fig. 4b), was gradually stronger than the asymptotic prediction within individual networks (colored line in Fig. 4b) and substantially better than single network when voxels were randomly sampled more than 1000. Of note, though several networks showed higher prediction performance than BAAS with limited voxels (e.g., n = 50), the prediction differences between networks and BAAS were not significant (Z scores ≤ 1.17). This analysis confirmed that whole-brain models exhibit significantly larger effect sizes than models using features from a single network. Moreover, consistent with previous studies on affective neural signatures^13,14^, the optimum BAAS performance was achieved when approximately 10,000 voxels were randomly sampled across the whole brain. Crucially, the performance was contingent upon a diverse selection of voxels derived from multiple brain systems. These findings further support the notion that information about affective arousal experiences is combined in the neural signature that extends across multiple systems.

In addition, we calculated the overlapping ratio between significant features of the BAAS and each large-scale functional network, subcortical network and cerebellar brain (Fig. 4c). The positive predictive weights of the model showed important overlaps with voxels of the visual (26.75%), default mode (16.72%), ventral attention (22.13%), and subcortical (10.83%) networks, and the negative one had a remarkable overlap with visual (31.37%), frontoparietal network (21.24%), somatomotor (16.67%) and dorsal attention (15.69%) networks. These findings further support the notion that arousal is represented across widely distributed brain networks.

## Affective arousal and autonomic arousal engage distinct neural representations in humans

Previous conceptualizations have hypothesized that affective arousal and autonomic arousal may operate through distinct neural mechanisms^19^. Given GSR has been validated as a stable and valid index of autonomic arousal^82^, we developed a whole-brain pattern for decoding autonomic arousal based on brain activation maps corresponding to various levels of GSR in study 25 (n = 25, leave-one-subject-out (LOSO) cross-validation procedure, see Supplementary Methods), consistent with the approach described by Taschereau-Dumouchel et al.^15^. The developed autonomic arousal signature accurately predicted the level of autonomic arousal (within-subject correlation coefficient r = 0.79 ± 0.03 SE; overall prediction-outcome correlation coefficient r = 0.61). To further validate the developed decoder, a two-alternative forced-choice test was applied in the Pavlovian threat conditioning task (study 24). In line with CS+ being accompanied by higher GSR than CS-, the autonomic arousal decoder response accurately classified CS+ versus CS- with 78 ± 5.5% SE accuracy (p = 3.01 ×10^05^, d = 0.91) (Fig. 5a). Of note, the autonomic arousal decoder developed in the current study used the same training dataset as that developed by Taschereau-Dumouchel et al.^15^. The only difference is that the current study employed a grey matter mask, whereas Taschereau-Dumouchel et al.^15^ used a whole-brain mask. As expected, the two decoders exhibit similar functional performance (the decoder from Taschereau-Dumouchel et al.^15^ classified CS+ versus CS- with 78 ± 5.5% accuracy (p = 3.01 ×10^05^, d = 0.87)) and spatial similarity (Pearson’s r across grey matter voxels is 0.92). Moreover, the generalizability of the autonomic arousal decoder was additionally validated in the original study using three independent datasets (for details see ref.^15^). Notably, in study 26, the autonomic arousal decoder also showed well generalizability (see below for detailed results).

**Fig. 5.**
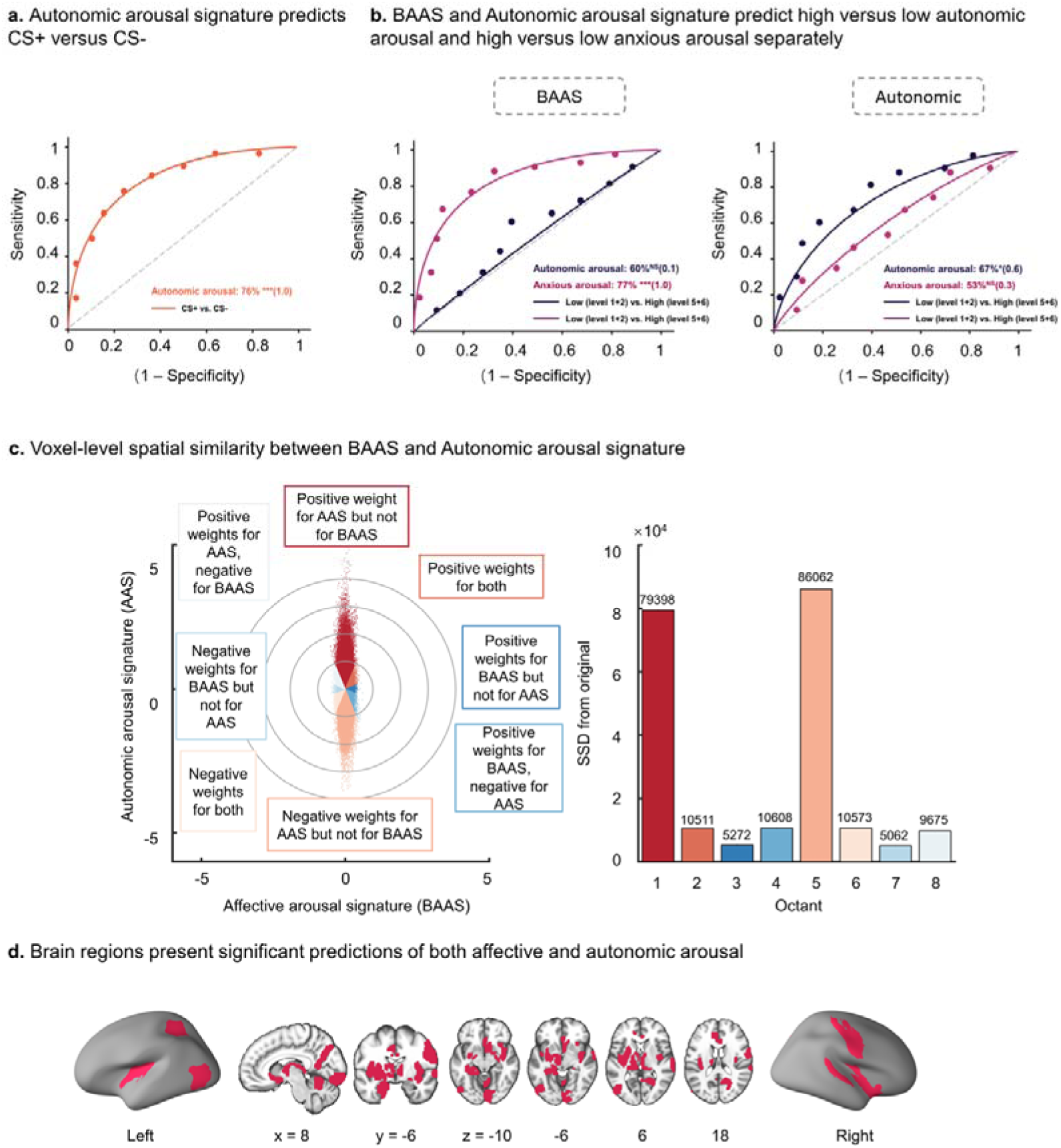
**Comparing neurofunctional decoders for affective arousal and autonomic arousal**. (**a**) During a Pavlovian threat conditioning fMRI task (study 24), the developed autonomic arousal signature could predict CS+ (associated with high GSR) versus CS- (associated with low GSR). (**b**) In study 26, BAAS more accurately predicts the anxious arousal experience during discrimination at high versus low intensity, while autonomic arousal more accurately predicts the autonomic arousal during discrimination at high versus low level. (**c**) Scatter plot displaying normalized voxel weights for affective arousal (BAAS, x-axis) and autonomic arousal (AAS, y-axis) signatures. Bars on the right side represent the sum of squared distances from the origin (0,0) for each Octant. Each Octant is assigned a different color, indicating voxels with shared positive or shared negative weights (Octants 2 and 6, respectively). Octant 1 represents selectively positive weights for the positive affect signature, Octant 3 represents selectively positive weights for the negative affect signature, Octant 5 represents selectively negative weights for the positive affect signature, Octant 7 represents selectively negative weights for the negative affect signature, and Octants 4 and 8 represent voxels with opposite weights for the two neural signatures. The numbers at the top of each bar indicate the number of voxels in each Octant. (**d**) Regions present a significant prediction of subjective affective arousal ratings and GSR. * P < 0.05, *** P < 0.001, NS not significant. BAAS, Brain affective arousal signature; GSR, Galvanic skin response; CS, conditioned stimulus.

To test whether affective arousal and autonomic arousal exhibit (partially) distinguishable neural representations, we employed a double dissociation approach which is a weaker form of the separate modifiability criterion that has been one of the main ways of dissociating mental processes in neuropsychological studies^83,84^. To this end, we first applied the BAAS and autonomic arousal signature to an independent fMRI dataset collected during a UVTA task from study 26^16^. This produced beta maps corresponding to anxious arousal ratings and categorized brain activation maps based on five different levels of GSR during anxious anticipation for each subject (see Supplementary Methods for details). We found that the BAAS significantly predicted anxious arousal ratings with r = 0.26 (p = 1.55×10LJLJ) and discriminated high (average of ratings 4 and 5) versus low anxious arousal experience (average of ratings 1 and 2) with high accuracy (77 ± 6.4%, p = 6.06 ×10LJLJ, d = 0.99). In contrast, the BAAS showed low predictive performance for GSR responses (r = 0.06, p = 0.41) and failed to classify high GSR (average of ratings 4 and 5) from low GSR (average of ratings 1 and 2) above chance level (60 ± 7.5%, p = 0.22, d = 0.05). Additionally, the autonomic arousal signature predicted autonomic arousal ratings with r = 0.21 (p = 1.81×10LJ3) and successfully classified high GSR (average of ratings 4 and 5) versus low GSR (average of ratings 1 and 2) with 67 ± 7.1% accuracy (p = 0.03, d = 0.61), but could not predict anxious arousal ratings (r = 0.08, p = 0.23) nor discriminate high anxious experience (average of ratings 4 and 5) from low anxious experience (average of ratings 1 and 2) (53 ± 7.6% accuracy, p = 0.76, d = 0.26).

To further test the hypothesis of partially separable neural representations underlying affective arousal and autonomic arousal, we explored the joint distribution of normalized (z-scored) voxel weights of these two patterns by plotting BAAS on the x-axis and the autonomic arousal signature on the y-axis (for similar approach see ref.^85^). As visualized in Fig. 5c, stronger weights across the whole brain (sum of squared distances to the origin [SSDO]) were located in the non-shared Octants (1 and 5, i.e., positive/negative weights for autonomic arousal signature but not for BAAS).

These dissociations would provide support for the notion that the two signatures do not exclusively rely on a single shared neurofunctional representation at the whole-brain level. However, it does not necessarily mean that affective arousal and autonomic arousal cannot be highly correlated at the same time nor that these two measurements exhibit distinct neural representations in each brain system. To further evaluate this aspect, we next tested whether some brain systems exhibit common representations of subjective affective arousal ratings and GSR on the local level. To this end, we retrained the two signature models by restricting their activation to each of 275 regions^75,76^ using a LOSO cross-validation procedure following Taschereau-Dumouchel et al. 2020^15^ and next predicted affective arousal ratings in study 1 and GSR in study 25 (see Methods for details). As shown in Fig. 5d. significant regions most commonly involved in both predictions include subcortical regions such as the amygdala, hippocampus, basal ganglia, and thalamus and cortical regions including the ACC, insula, precentral gyrus, and superior parietal lobule as well as some cerebellar regions.

Together, these results suggest that the neural representations of affective arousal are distinguishable from those of autonomic arousal on the whole-brain level, while common brain representations may underlie both affective and autonomic arousal processes at the local level.

## BAAS improves the specificity of previously established affective neural signatures

To examine whether the BAAS could enhance the specificity of previously developed affective neural signatures, we conducted a series of analyses with two signatures. The visually negative affect signatures (VNAS) was designed specifically to respond to visually induced negative affect without encompassing other forms of negative experience or more positive experiences^34^. In study 1, VNAS was capable of distinguishing negative (i.e., fear and disgust) from positive (i.e., happiness), and neutral stimuli with 76.19% and 93.65% accuracy, respectively (Fig. 6a and Table S4; for detailed descriptions of each emotional video category in study 1 see Supplementary Methods). Despite these high accuracies, the specificity of the VNAS was limited due to higher response to positive compared to neutral videos (accuracy = 75.40%, p = 9.61 ×10^-9^). We assessed whether incorporating arousal as control could enhance the prediction specificity of the VNAS. By adjusting for arousal rating, we found that the predictions for negative affect were maintained, with VNAS effectively distinguishing negative from positive and neutral videos, achieving 67.46% and 61.91% accuracy, respectively. Importantly, post-adjustment, the VNAS showed comparable responses to positive and neutral videos (accuracy = 51.59%, p = 0.79). These findings demonstrated the potential of arousal rating regression in refining the specificity of the VNAS. Crucially, we found that adjusting the responses of the VNAS based on BAAS responses could improve prediction specificity to comparable levels, if not identical levels, to those achieved through direct arousal rating adjustments (Fig. 6a; see also Table S4). Following correction, VNAS showed comparable responses to positive and neutral videos (accuracy = 52.38%, p = 0.66). Notably, the adjusted approaches also allowed the VNAS to predict negative versus positive videos more accurately than negative versus neutral videos, underscoring the utility of controlling arousal in refining prediction specificity. These findings were further replicated in study 17, where positive, negative, and neutral emotions were elicited by pictures without acquiring arousal ratings.

**Fig. 6.**
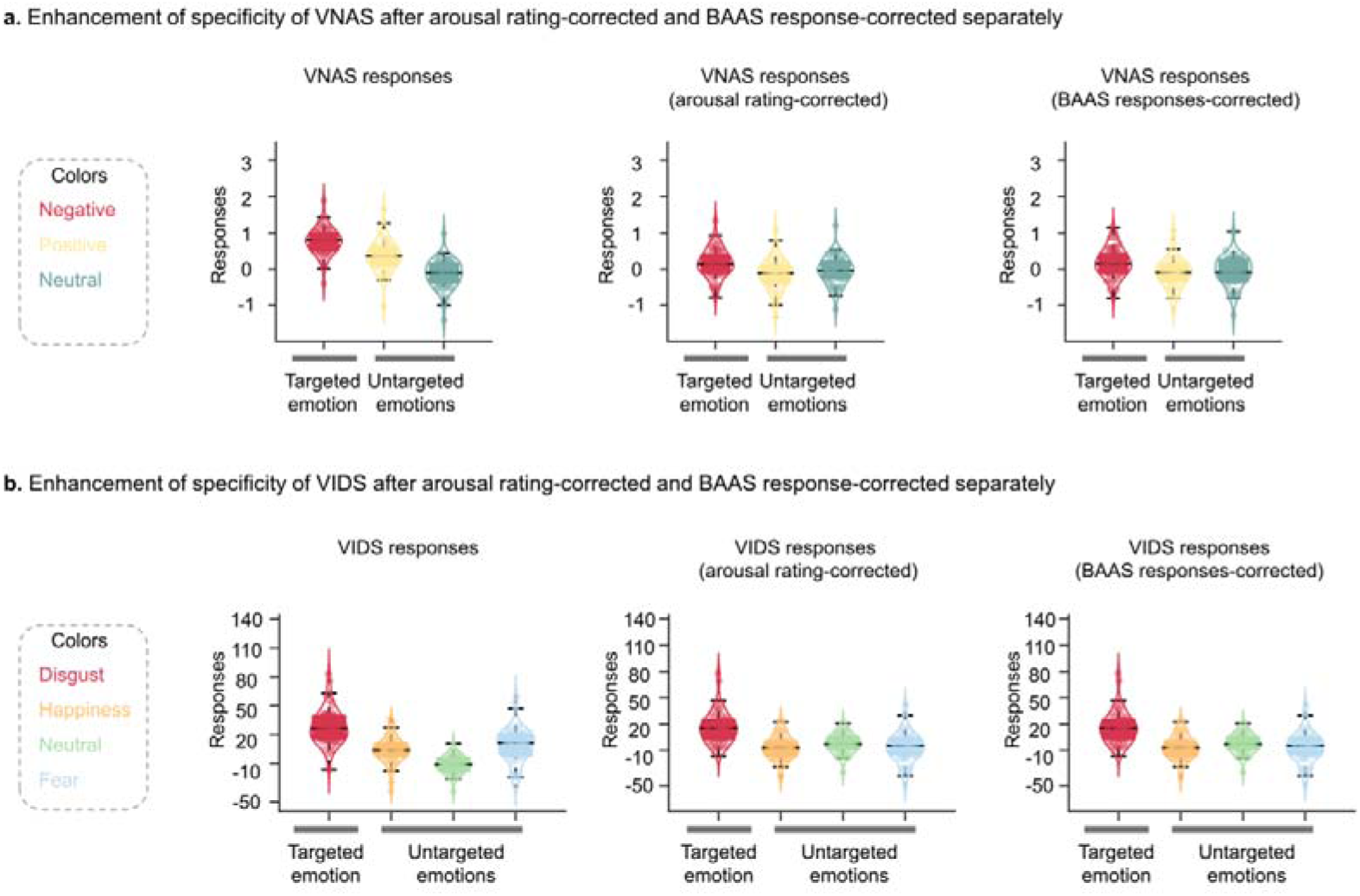
The application of BAAS enhances the specificity of VANS and VIDS in study 1. (**a**), in the left panel, VNAS displayed higher responses to negative affect, while it also significantly responded to positive compared to neutral conditions; in the middle and right panels, after two correction methods (actual arousal ratings, BAAS response correction), VNAS showed similar responses to positive and neutral conditions. For details, see Supplementary Results. (**b**), in the left panel, VIDS showed higher responses to targeted emotion (i.e., disgust), while it significantly responded to happiness and fear compared to neutral emotions; in the middle and right panels, after arousal-rating-corrected or BAAS-responses corrected, VIDS still performed higher response to disgust, while it showed comparable responses to happiness, fear, and neutral emotions. VNAS, visually negative affect signatures; VIDS, visually induced disgust signature.

Results consistently showed that regressing out the BAAS response enhanced the prediction specificity of the VNAS (see Supplementary Results for statistic details).

To further examine the reliability of our approach, we replicated the procedures using another affective neural signature, i.e., visually-induced disgust signature (VIDS)^13^. In study 1, VIDS predicted subjective disgust with robust discriminative power in distinguishing disgust from fear, happiness, and neutral videos, achieving 71% to 97% accuracies (Fig. 6b and Table S5). However, the VIDS also showed a heightened response to happiness (accuracy = 81.75%, p < 1.00×10^-10^) and fear (accuracy = 83.33%, p < 1.00 ×10^-10^) compared to neutral videos. Additionally, the VIDS exhibited a higher response to fear compared to happiness videos (accuracy = 64.29%, p = 1.70× 10^-3^). To address this issue, we incorporated adjustments for arousal ratings and BAAS responses. Following these adjustments, VIDS showed comparable responses to happiness and neutral videos, fear and neutral videos as well as fear and happiness videos (see Fig.6b and Table S5).

Collectively, our findings demonstrate the substantial potential of BAAS in refining the specificity of affective neural signatures, highlighting its prospective utility in advancing future models of affect prediction.

## Discussion

Arousal represents a key dimension of the affective space and accompanies all affective experiences^86^. Despite its central role in overarching models of emotion and extensive animal models focusing on the autonomic or vigilance components of arousal a comprehensive and accurate neurofunctional model of the conscious affective experience component is lacking. Establishing such a model may allow to test whether arousal is represented independent of valence, separable from the representation of autonomic arousal and may facilitate accurate neural decoding of specific emotional states^47,49^. We combined multisensory naturalistic neuroimaging with predictive models to develop and extensively evaluate a sensitive and generalizable brain signature for affective arousal (BAAS). The signature accurately predicted affective arousal on the group and individual level in the discovery cohort and showed robust predictive performance in an independent replication sample that covered a different emotional space. Moreover, extensive evaluation of the BAAS across a series of independent neuroimaging datasets confirmed the efficacy of the BAAS in predicting affective arousal across valence domains, modalities (e.g., visual, auditory, sensory, and noxious stimulation), experimental paradigms, and samples, as well as the high specificity of the BAAS in capturing affective arousal component. To determine the core systems of affective arousal in the human brain, we combined diverse multivariate and univariate analyses. The biological plausibility of the BAAS was further validated by shared common features between the BAAS and conjunction meta-analytic maps of both positive and negative affect, as derived from Neurosynth, which together with the topographic distribution of the most predictive regions of the BAAS, indicates that the amygdala, parahippocampus, PAG, and medial prefrontal systems contribute to high affective arousal across valence domains. Our work also evidenced that the neural representation and predictive domain of the BAAS were moreover separable from a signature predictive of autonomic arousal, indicating that the conscious emotional experience and the autonomic components are represented in partly dissociable neural representations (in line with refs.^15,16^). Last, and in line with the ‘overshadowing’ concept of high arousal overshadowing the determination of specific emotional states, we demonstrated that the BAAS could be successfully employed to considerably enhance the specificity of existing neural signatures for the target emotional state, suggesting a high application potential for the generation of more specific neuromarkers for mental processes. Together, our findings demonstrate that affective arousal can be accurately decoded from distributed brain systems encompassing both subcortical and cortical areas, with the predictive accuracy of no single region or network surpassing that of the comprehensive whole-brain model. This underscores the complexity and integrative nature of neural mechanisms underlying affective arousal^39,77^. We provide the first comprehensive neurofunctional representations of affective arousal and demonstrate that the corresponding distributed neural representation cuts across the valence domain is separable from autonomic arousal and overshadows precise decoding of specific emotional states.

The current findings revealed that affective arousal depends on the coordinated engagement of distributed representations throughout the entire brain rather than on activity in isolated brain systems, which is consistent with recent neural decoding studies demonstrating that affective processes are represented across brain systems^12–14,87^. Specifically, affective arousal experience requires concerted engagement of brain-wide distributed representations with comparable strong contributions of subcortical regions involved in affective, autonomic and motor responses to highly salient stimuli in the environment (amygdala, parahippocampus, PAG, and caudate)^30,88–90^, anterior cortical midline structures associated with global arousal and implicit emotion regulation (medial prefrontal cortex, ACC)^74,91,92^, anterior insular systems involved in interoceptive and affective awareness^93,94^ and sensorimotor integration (occipital cortex, SMA)^95^, as well as regions that have been identified as key mediators of wakefulness and alertness in animal models (brainstem)^19,25^. Furthermore, no individual large-scale brain network significantly outperformed the BAAS in predictive performance (see Fig. 4b). Conjunction analysis revealed that the core systems (thresholded BAAS) of affective arousal are distributed across several large-scale brain networks. Notably, the DMN, ventral attention network and subcortical network exhibited significant proportions of positive predictive weights for BAAS. These findings align with previous literature suggesting that the DMN (e.g., anterior medial prefrontal cortex, portions of the ventromedial prefrontal cortex) and salience network (e.g., ACC, insula, amygdala) play crucial roles in affective arousal and conscious emotional experience^19,29,96^. From an evolutionary perspective, the engagement of these systems may facilitate the detection of salient information in the environment and trigger autonomic processes to generate the adaptive cognitive and homeostatic response^78^. Together, these findings underscore that distributed representations of affective arousal support appraisal^97^ and constructionist^77^ theories of emotions that propose the shared but also distinct distributed neural systems that span cortical and subcortical regions facilitate subjective affective experiences.

From a larger conceptual perspective, affective neuroscience and affective space theories propose that high affective and autonomic arousal will cut across the valence domain and characterize high-intensity positive and negative emotional experiences. In support of these conceptualizations, previous imaging meta-analysis uncovered overlapping brain regions (e.g., amygdala, striatum, thalamus, medial prefrontal cortex, ACC, and SMA) implicated in both positive and negative emotions^28^, which may represent systems involved in general emotion processing (e.g. appraisal) as well as affective and autonomic arousal^5,9^. Testing the generalization of the BAAS revealed that the signature robustly decoded high positive and negative experiences to a similar extent but did not differentiate between them. Together with a flanking meta-analytic strategy identifying overlapping regions between the BAAS and both positive and negative emotional experiences, these findings indicate that the amygdala, parahippocampal gyrus, inferior and dorsomedial frontal regions code affective arousal irrespective of valence (for supporting evidence from conventional strategies see^27,28,98–100^). To further differentiate the contribution of hard-wired physiological arousal responses to the valence-independent affective arousal signature we employed a validated autonomic arousal signature and demonstrate that the BAAS has a considerably higher specificity for affective arousal. These findings are in line with a growing literature suggesting that the neural representations of the hard-wired autonomic responses and the conscious affective experience are distinguishable (for example, refs.^15,16^) and underscore that the BAAS captures the affective component. Notably, consistent with previous research^101^, we found that – at the local level – a number of subcortical (amygdala, thalamus), cerebellar and insular-ACC regions predicted both affective and autonomic arousal, emphasizing the critical role of these systems in general arousal processes. Our work thus presents converging evidence demonstrating that while distributed neural representations constitute distinguishable substrates of different arousal facets (i.e., affective and autonomic arousal), there are some regions that encode affective and autonomic arousal with similar neural representations.

As outlined in different domains, including subjective emotional experiences, emotion recognition as well as neural and autonomic signatures, increasing arousal progressively overshadows the characteristics of specific emotional states. From the perspective of affective neuroimaging, it is therefore critical to control for non-specific emotional processes such as arousal which is inherently associated with the affective experience^11,47,49^. We explored this conceptual and technical perspective by employing the BAAS in combination with evaluated robust signatures for general negative affect (VNAS^34^) and disgust (VIDS^13^). In line with the overshadowing concept, the signatures showed limited specificity during immersive high-arousing emotion processing (study 1, study 17 for convergent evidence see also refs.^35,47^). While the arousal contribution considerably affects neural decoding as well as conventional affective neuroimaging experiments^102^ solutions to this issue have not yet been effectively identified. We here capitalized on the BAAS to critically enhance the predictive specificity of these affective neural signatures by directly regressing out the experience of arousal. The enhanced specificity demonstrated the effectiveness of controlling for the influence of affective arousal on the prediction of emotional states. Furthermore, our findings indicate that adjusting the BAAS response yields a comparable effect to adjusting actual subjective arousal ratings. This holds particular importance given that arousal ratings are not consistently gathered in affective neuroimaging research (or the assessment can strongly interfere with the actual mental process under investigation). Our work provides a simple yet potent strategy by employing the BAAS to mitigate the impact of affective arousal, thereby significantly improving the predictive specificity of other affective neural signatures.

While the current study shows that the BAAS generalizes to multimodal affective arousal, it does not mean that each feature (i.e., voxel) contributes to the predictions of the BAAS similarly across stimulus types. Previous studies have shown that there are common and stimulus-type-specific brain representations of affective experience^1,34^. Future work is necessary to understand the stimulus-type-specific neural representations of the affective arousal domain.

Additionally, some important brain systems of affective arousal are anatomically and functionally connected (e.g., amygdala and hippocampus) and some previous studies demonstrated the engagement of brain networks in wakefulness arousal fluctuations (for example, ref ^103^), underscoring a potential role of brain pathways and network level communication in affective arousal^88,104^ and future research may aim to deepen our understanding of the neurobiological basis of affective arousal by mapping the brain pathways that underlie this process. Moreover, the current study primarily focused on elucidating the functional brain architecture underlying general affective arousal. However, specific neural representations may correlate strongly with self-reported arousal for either positive or negative events. Future studies could investigate the distinct neural patterns underlying the intensity of positive and negative affective experiences. Lastly, considering the characteristic of affective arousal, BAAS was mainly tested in sympathetic arousal-related processes; distinguishing the contributions of parasympathetic and sympathetic arousal to these neural patterns may provide valuable insights into the regulatory mechanisms of affective states in future studies.

In summary, we established a comprehensive distributed, generalizable, and specific predictive neural signature of affective arousal. The signature was validated and generalized across modalities, valence, paradigms, participants, and MRI systems. We showed that the neural representations of affective arousal are encoded in multiple distributed (large-scale) brain systems rather than isolated brain regions. The affective arousal signature cut across the valence domain and was distinguishable from hard-wired autonomic arousal, indicating a valence-unspecific, yet affective arousal-specific neural representation. Based on the characterization of the BAAS, we further developed a novel approach to enhancing the specificity of affective neural signatures. The current work extends our understanding of the neurobiological underpinnings of affective arousal experience and provides objective neurobiological measures of affective arousal. These measures can complement self-reported affective arousal and serve as effective supplementary tools to enhance the prediction specificity of affective models.

## Methods Overview

This research utilized data from n = 26 studies. Study 1 served as the primary study and discovery cohort, which was employed for the main analysis and development of the affective arousal signature (i.e., BAAS). Study 2 involved novel emotional video stimuli to validate the identified affective arousal signatures in an independent cohort. Studies 3 to 18 comprised multiple stimuli and modalities and were utilized for the evaluation of generalizability. Studies 19 to 24 involving cognitive processes were applied to test the effect of potential confounding factors in the model. The autonomic arousal signature was developed in study 25 and validated in study 24. Finally, the affective arousal signature and autonomic arousal signature were compared using data from study 26. Additional details for each study are provided in Table S1, and an overview of the main analysis is depicted in Fig. 1.

## Participants in study 1

Sixty-three healthy, right-handed participants were recruited from the University of Electronic Science and Technology of China (UESTC) to undergo fMRI scans during a subjective arousal rating paradigm using movie clips. All participants had no current and past physical, neurological, or psychiatric disorders and had not taken any medication in the month before the experiment. Three participants were excluded due to a lack of high arousal experience (the number of high arousal clips was less than 2, where high arousal was defined as an arousal rating score above 6), resulting in a final analysis of n = 60 participants (23 females; mean ± SD age = 19.55 ± 1.87 years).

The experiment procedure was approved by the local ethics committee at the University of Electronic Science and Technology of China and was conducted under the latest version of the Declaration of Helsinki. Before the regular experiments, all participants were informed that they would be shown fearful and disgusting movie clips on the screen and completed written informed consent. After the experiment, participants received monetary compensation for study participation.

## Movie stimuli, arousal rating paradigm, and procedure in study 1

The movie stimuli were selected from a large pool of clips that were downloaded from an online sharing platform. This selection process was performed by two research assistants. A total of 80 movie clips, each lasts ∼25s and were categorized as positive, negative, and neutral. During the pre-study, independent participants (n = 26; 10 females, age = 21.62 ± 1.92) were asked to attentively watch the clips, and rate their level of arousal, valence, and emotional consistency using a 9-point Likert scale (1 = low arousal or negative or low consistency, 9 = high arousal or positive or high consistency), and select one emotional word (such as ‘fearful’, ‘disgusting’, ‘angry’, ‘sad’, ‘surprising’, ‘happy’, ‘neural’, or ‘ambiguous’) that could accurately describe their emotional experience following each clip. The stimuli rating experiment was delivered using PsychoPy^105^ version 1.85 under Windows. Low arousing neutral and high arousing positive and negative videos were selected as fMRI experiment stimuli while high emotional consistency (rating > 7) was included to serve as the reference condition.

The current study employed a within-subject design. In the arousal rating fMRI paradigm (see Fig. 1a), participants were presented with 40 arousing movie clips over 3 runs. Each run included 14 or 12 clips, which depicted humans, animals, and scenes. The order of runs and trials within a run was the same for all participants, and the order of conditions was counterbalanced before the experiment. Each video lasted approximately 25s and was followed by a white fixation cross on a black background for 1-1.2s. During the subsequent 5s period, participants were instructed to report their level of arousal during each movie using a 9-point Likert scale, where 1 indicates very low arousal and 9 indicates very high arousal. This was followed by a 10-12s wash-out period. Participants watched a total of 20 low-arousal neural movie clips, 10 high-arousal positive clips, and 10 high-arousal negative clips (details provided in Table S2). Stimulus presentation and behavioral data acquisition were controlled using MATLAB 2014a (Mathworks, Natick, MA) and Psychtoolbox (http://psychtoolbox.org/).

Before the fMRI experiment, all participants were trained on the laptop to understand the definition of ‘arousal’. After the scanning, they were asked to select the correct screenshot of movie clips presented during the fMRI experiments in a post-test.

## fMRI data acquisition in study 1

fMRI data were collected using standard sequences on a 3.0-Tesla GE Discovery MR750 system (General Electric Medical System, Milwaukee, WI, USA). Structural images were acquired by using a T1-weighted MPRAGE sequence (TR = 6ms; TE = 2ms, flip angle = 9°, field of view = 256 × 256 mm^2^; matrix size = 256 × 256; voxel size = 1 × 1 × 1 mm). Functional images were acquired using a T2*-weighted echo-planar sequence (TR = 2000 ms; TE = 30ms; slices = 39; flip angle = 90°; field of view = 240 × 240 mm; matrix size = 64 × 64, voxel size: 3.75 × 3.75 × 4 mm).

## Participants in study 2

A total of 38 healthy right-handed UESTC students were initially recruited for the study. Following exclusion criteria based on low arousal ratings for positive or negative videos in the fMRI experiments, 36 subjects (17 female, mean ± SD age = 21.03 ± 2.22) were enrolled in the final analysis. All participants were free of current and past physical, neurological, or psychiatric disorders and had no contraindications for MRI. Provided written informed consent before participation and the experimental procedures were approved by the local ethics committee at UESTC, in accordance with the latest version of the Declaration of Helsinki. Monetary compensation was provided to all participants for their participation in the study.

## Movie stimuli, arousal rating paradigm in study 2

Stimuli used in study 2 were entirely different from those used in study 1. All movie clips (∼60s duration of each) were selected by two researchers to ensure a diverse and non-overlapping set of stimuli. An independent sample of 52 participants (32 females, mean ± SD age = 21.00 ± 2.51) reported the valence, arousal, emotional consistency, and emotional experience (‘fearful’, ‘disgusting’, ‘sad’, ‘ambiguous negative’, ‘happy’, ‘exciting’, ‘romantic’, ‘ambiguous positive’, and ‘neutral’) while watching each video. Of note, when participants had multiple positive or negative emotions, the labels ‘ambiguous positive’ or ‘ambiguous negative’ were chosen (e.g., if a video evokes both happiness and excitement). The selected fMRI video stimuli were chosen based on the results of the behavioral experiment as (1) they were rated high emotional consistency (rating > 7) and (2) their core emotional experience was accurately described (i.e., no positive videos labeled with negative emotions, and no negative videos labeled with positive emotions).

In the arousal rating fMRI paradigm, there were a total of 28 one-minute videos. These videos represented two affective states (9 positive clips and 9 negative clips) as well as a neutral state (10 neutral clips). More details can be found in Table S2. During fMRI, all stimuli were presented in three runs, with either three or four clips from each category in each run. The fMRI paradigm used in the validation cohort was the same as the one used in the discovery cohort, except for differences in the presentation time settings for the stimuli.

## fMRI data acquisition in study 2

fMRI data were collected on a 3.0-Tesla GE Discovery MR750 system (General Electric Medical System, Milwaukee, WI, USA). Functional images were acquired using a T2*-weighted echo-planar sequence (TR = 2000ms; TE = 30ms; slices = 36; flip angle = 90°; field of view = 200 × 200 mm^2^; matrix size = 64 × 64, voxel size: 3.125 × 3.125 × 3.8 mm). To improve spatial normalization and exclude subjects with apparent brain pathologies structural images were acquired by using a T1-weighted MPRAGE sequence (TR = 8ms; TE =3ms, flip angle = 8°, field of view = 256 × 256 mm; matrix size = 256 × 256; voxel size = 1 × 1 × 1 mm).

## fMRI data preprocessing in studies 1 and 2

Before the automated preprocessing, the first 4 volumes of each run were removed to stabilize image intensity. Imaging data were preprocessed using FMRIPREP 21.0.0(RRID: SCR_016216; https://github.com/nipreps/fmriprep)^106^, a Nipype 1.6.156 based tool that integrates preprocessing routines from different software packages, including slice time corrected (using 3dTshif from AFNI), motion correction (using mcflirt from FSL v6.0.5.1), brain tissue segmentation of cerebrospinal fluid (CSF), white matter, and gray matter (using FAST from FSL v6.0.5.1), coregistration (using boundary-Based registration with six degrees of freedom from FLIRT), and affine transformation of the functional volumes to the 2 mm isotropic T1w anatomical and subsequently to MNI space (using ANTs v2.3.3). To perform denoising, we fit a voxel-wise general linear model (GLM) for each participant. Specifically, in line with the ref^107^, we removed variance associated with the mean, linear and quadratic trends, the average signal within anatomically-derived eroded CSF mask, the effects of motion estimated during the head-motion correction using an expanded set of 24 motion parameters (six realignment parameters, their squares, their derivatives, and their squared derivatives) and motion spikes (FMRIPREP default: framewise displacement (FD) > 0.5mm or standardized DVARS > 1.5). For arousal rating data we additionally constructed a task-related regressor with rating period convolved with canonical hemodynamic response function (HRF).

## Data averaging

To develop the subjective affective arousal signatures, we first generated arousal-related activation features for each subject. For each participant, we concatenated the fMRI data with the same arousal level rating during the stimuli presentation period. Notably, the onset time for each video was shifted by 2-3 TRs due to the hemodynamic lag. Next, we calculated the voxel-wise mean activation for each arousal level and each subject separately. For the single-trial analysis, we also acquired the fMRI data for each stimulus. The same method was employed for the fMRI data collected in study 2.

## Brain affective arousal signature development and evaluation

In study 1, we developed a predictive model of arousal ratings based on whole-brain activation (i.e., BAAS). Whole-brain data mask with gray matter from this study was used. We employed the support vector regression (SVR) algorithm using a linear kernel (C=1) implemented in the Spider toolbox (http://people.kyb.tuebingen.mpg.de/spider) with voxel-wise activation maps from each rating as the predictor (true arousal ratings as outcome). We employed a 10×10-fold cross-validation procedure to evaluate the identified signature, which involved random assignment of all participants to 10 subsamples, comprising 6 individuals, using MATLAB’s cvpartition function. The optimal hyperplane was then derived from the multivariate pattern observed in the training set, consisting of 54 participants, and subsequently evaluated using the excluded 6 participants in the test set. This rigorous procedure was repeated ten times, with each subsample serving as the testing set once. To thoroughly evaluate the neurofunctional signature, we employed a range of robust metrics including prediction-outcome correlation, the average root mean squared error (RMSE), and the coefficient of determination (R^2^). Specifically, we estimated the Pearson correlation (r) between cross-validation predicted and observed ratings (that is, true arousal ratings) to indicate the effect sizes, including overall (between and within-subject; 396 pairs in total) and within-subject Pearson correlation coefficient. RMSE illustrated the overall prediction errors, while R^2^ represented the percentage of arousal rating variance accounted for by the prediction models. The formula is:

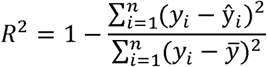

where y_i_ is the true arousal rating for the i-th sample, 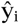 is the model predicted rating for the i-th sample, and 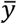 is the mean of all arousal ratings.

To further evaluate the performance of the arousal-predictive pattern (trained on the discovery cohort), we applied BAAS to validation and generalization datasets. In the validation cohort (study 2), signature responses were calculated by taking the dot product of the BAAS weight map with the test images for each participant for each arousal level. The overall and within-subject Pearson correlation coefficient, RMSE, and R^2^ between the predicted and observed ratings were used to determine the performance. Furthermore, we evaluated the performance of BAAS in predicting different levels of arousal experience in the validation cohort by assessing classification accuracies using a forced-choice test. The intensity of affective arousal was classified into three levels: low intensity (1-3 scores), medium intensity (4-6 scores), and high intensity (7-9 scores). We compared the distinctive responses under two different conditions for each level of intensity (low versus medium, medium versus high, low versus high), and considered the condition with the higher signature response as being more associated with increased affective arousal experience.

## Within-subject trial-wise prediction

We aim to test whether the BAAS could effectively predict individual trial-by-trial subjective arousal levels. To this end, we performed a single-trial analysis^13,14^, which was achieved by constructing a GLM design matrix with separate regressors for each stimulus. Of note, for each subject we did not exclude any trials from subsequent analyses. Next, we calculated the BAAS pattern expressions of single-trial voxel-wise activation maps, which involved taking the dot-product of vectorized activation images with the BAAS weights. We then correlated the pattern expressions with the true ratings separately for each participant. The statistical significance was evaluated by prediction-outcome Pearson correlation for each subject separately. For the discovery cohort, consistent with prior studies^13,14^, we used the 10×10-fold cross-validation procedure to obtain the BAAS response of each single-trial activation map for each subject. We additionally calculated the within-subject trial-wise prediction of BAAS in the validation cohort using the model trained on the discovery cohort (i.e., the BAAS).

## Generalization of BAAS performance

To evaluate the generalizability of the affective arousal signature, we examined its response in sixteen independent datasets from opening research (details see Table S1). In studies 3-6, we assessed classification accuracies of BAAS between high and low arousal feelings induced by various negative stimuli and modalities using a two-alternative forced-choice method based on the pattern expressions (i.e., the dot-product of BAAS weight map and test image plus the intercept).

In studies 7-16, we conducted a one-sample t-test to examine whether there existed significant differences between BAAS responses (via dot-products) for positive and baseline contrasts, allowing us to assess the robustness of the BAAS in positive contexts. Study 17 (a large sample of participants received positive, negative, and neutral image stimuli) was used to examine the discriminability of BAAS during positive and negative conditions within the same task, further validating its generalization across different affective states. Study 18 was chosen to test BAAS response in a mental imagery task without direct external stimuli. This provided evidence of the signature’s performance in conditions where affective arousal is internally generated rather than externally induced.

## Specificity of BAAS performance

To estimate the specificity of the affective arousal signature, in the validation cohort (study 2) and studies 19-24, we conducted a series of two-alternative forced-choice tests based on pattern expressions and one-sample t-tests based on BAAS responses. Study 2 was used to test whether the BAAS was specific to arousal and not merely a reflection of valence. Studies 19 and 20 included incongruent versus congruent trials in Eriksen Flanker and Simon tasks to assess potential confounding factors such as attention-grabbing. Studies 21 and 22, involving a comparison of 1-back versus 0-back trials in the N-back task and remembered positive/negative items versus remembered neutral items in the emotional memory task, were used to evaluate potential confounding factors such as memory. Studies 23 and 24 were chosen to test whether this affective arousal signature showed limited predictive performance to wakefulness and autonomic arousal.

## Identifying neurofunctional characterization of affective arousal

We performed a comprehensive analysis to identify the core brain regions (i.e., having voxels that reliably contribute to model weights) associated with subjective affective arousal processes. To this end, we conducted a bootstrap test using 5000 samples with replacements from the discovery cohort to determine the voxels that made the most reliable contribution to the prediction and calculated Z-scores at each voxel with the mean and standard deviation of the sampling distribution. The resulting maps were thresholded voxel-wise at q < 0.05 (FDR corrected). Additionally, we performed parcellation -based (274 regions from the Brainnetome atlas^75^, and the PAG from Roy et al^76^) prediction analysis to identify local regions predictive of affective arousal in both discovery and validation cohorts. Specifically, in each region, we used 10×10-fold cross-validation within the discovery cohort in study 1 and applied the brain model (i.e., BAAS) to new individuals in the validation cohort (study 2). We then conducted the conjunction analysis to identify brain regions with significant (p < 0.001 uncorrected, equivalent to FDR corrected q < 0.002) and consistent performances in two cohorts.

To evaluate the validity and neurobiological plausibility of core systems, we additionally employed the following analyses to identify the specific neural basis underlying the subjective experience of affective arousal: (1) consistent with previous studies^13,14^, we performed multivariate analyses to locate brain regions that predictive of arousal ratings. To this end, we first ran a separate prediction analysis (linear SVR with CLJ=LJ1) for each participant in the discovery cohort (study 1) using their single-trial activation maps. A one-sample t-test across these individualized predictive models was performed to evaluate the consistency of each weight for every voxel in the brain (qLJ<LJ0.05, FDR corrected; see Supplementary Methods for detailed methods). (2) We obtained first-level univariate parametric modulation beta maps and performed a one-sample t-test on all images to identify regions of activation associated with arousal ratings. (3) We compared the BAAS with the conjunction of ‘positive affect’ and ‘negative affect’ Neurosynth maps (association test q < 0.05, FDR corrected; ‘positive affect’ studies n = 1,095; ‘negative affect’ studies n = 769) to confirm the biological plausibility of core systems. Overall, these analyses provide an extensive assessment of the neurobiological basis of the subjective experience of affective arousal.

To test whether affective arousal processing, as a component of ‘core affect’, could be characterized by activations within a single network or if it is neurobiologically supported by large-scale networks, as proposed by psychological construction theories^28^, we compared the prediction performances of large-scale networks against whole brain approaches. The networks of interest included seven resting-state functional networks, a subcortical network, including the amygdala, hippocampus, striatum, and thalamus, as well as the cerebellar brain (from Brainnetome atlas^75^). To conduct this analysis, we selected and tested the model for the network using discovery data with a 10×10-fold cross-validation procedure. two-tailed Z-test was used to examine the differences between predictive performances.

As the experience of affective arousal was distributed across multiple neural systems and supported by large-scale networks (see results), we created a polar plot to illustrate the proportion of overlaps between core systems (thresholded BAAS) against a set of anatomical parcellations (seven resting-state functional networks, one subcortical network, and cerebellar brain). To reduce potential biases arising from different atlases, we continued to use the Brainnetome atlas^75^.

## Comparing the affective arousal signature and autonomic arousal signature

Previous literature has highlighted the potential distinction between affective arousal and autonomic arousal^19^. To compare these, we trained a whole-brain autonomic arousal decoder in the study 25 (n = 25). Detailed information about the data and training methods for study 25 is provided in the Supplementary Methods. This decoder was also applied to study 24 (n = 58), where we assessed the two-alternative forced-choice classification accuracy between CS+ (associated with high GSR) and CS- (associated with low GSR) to validate the reliability of the identified autonomic arousal signature. Subsequently, the differences between the BAAS and the autonomic arousal signature were determined by: (1) calculating the two-alternative forced-choice classification accuracies between different anxious arousal intensity levels (as well as GSR) based on pattern expression in study 26 (details of this study see Supplementary Methods), and (2) plotting the joint distribution of normalized whole-brain weights of affective arousal and autonomic arousal patterns (see Supplementary Methods for details). Finally, we conducted an exploratory conjunction analysis to identify the brain regions involved in the significant prediction of both affective arousal ratings and GSR. To this end, we retrained affective arousal and autonomic arousal decoders within each of the selected 274 regions based on the Brainnetome atlas^75^ and the PAG from Roy et al.^76^ (n = 275 regions in total; https://osf.io/m3fjb/) using LOSO cross-validation procedure to predict arousal ratings in study 1 and GSR in study 25. This provided us with a correlation coefficient between the predicted and true values for each decoder, within each region. After multiple comparisons correction, we conducted an exploratory conjunction analysis to determine brain regions predictive of both affective arousal ratings and GSR (q < 0.05, FDR corrected).

## Evaluating the potential of BAAS in enhancing the specificity of affective neural signatures

Previous studies have developed and evaluated whole-brain affective decoders for general negative affect experience (VNAS^34^, from another lab) and subjective disgust experience (VIDS^13^, from our lab). To evaluate the potential of BAAS in improving the specificity of affective neural signatures, we conducted a series of analyses. Firstly, VNAS were applied to evaluate responses to a collection of emotional videos, including negative (i.e., fear and disgust), positive (i.e., happiness) and neutral emotions in study 1. We quantified the pattern responses (i.e., the dot-product of model and test brain activation map) and assessed the predictive accuracies using a single-interval classification. This method for estimating neural response patterns was consistently applied in subsequent analyses. Secondly, we examined whether controlling for arousal ratings in VNAS predictions could enhance specificity. Specifically, we used BAAS to predict the arousal scores of each subject during each category of emotional experience. After regressing out the predicted arousal scores from VNAS responses across all conditions, we reevaluated the predictive performances of the VNAS under each condition. Finally, we examined the predictive accuracy of VNAS after adjusting their responses based on the BAAS responses. This involved obtaining the BAAS response for each condition and then controlling for these BAAS responses in the VNAS responses across all conditions. We also validated these findings in study 17. Specifically, we tested testing whether adjusting the VNAS responses based on BAAS responses could enhance the predictive specificity of the VNAS when the dataset did not include any arousal ratings. To further validate our findings, we employed another affective neural signature, i.e., VIDS. In study 1, VIDS was used to predict responses to disgust, fear, happiness, and neutral videos, following the same analysis pipeline as described above.

## Behavioral data analysis

We used MATLAB script to check all behavioral data and exclude the participants who had a low affective experience. Statistical analyses of behavioral data were carried out using SPSS software (version 22; IBM Corp., Armonk, NY). We examined whether participants had strong arousal during the stimuli presentation (details see results).

## Acknowledgments

We thank J. He, D. Liu, and K. Fu for assisting in data collection; M. Čeko and colleagues for sharing data included in study 4; T. Wager and colleagues for sharing data included in study 5; X. Liu and colleagues for sharing data included in study 6 and study 26; P. Kragel and colleagues for sharing data included in studies 7 – 16; D. Coynel, D. J.-F. de Quervain, and colleagues for sharing data included in study 17; S. Lee and colleagues for sharing data included in study 18; A.C. Kelly, and colleagues for sharing data included in study 19 and 20; X. Xu and colleagues for sharing data included in study 21; Y. Gu and colleagues for sharing data included in study 23; V. Taschereau-Dumouchel and colleagues for sharing data included in study 25. Any opinions, findings, conclusions, or recommendations expressed in this publication do not reflect the views of the Government of the Hong Kong Special Administrative Region or the Innovation and Technology Commission. This work was partly supported by the National Natural Science Foundation of China (grant nos. 32250610208 and 82271583 to B.B., and 32300862 to F.Z.), National Key Research and Development Program of China (grant no. 2018YFA0701400) to B.B., the Natural Science of Foundation of Chongqing (CSTB2023NSCQ-MSX0889) to F.Z., Chongqing Talent Program (2024YC035) to F.Z., Fundamental Research Funds for the Central Universities (SWU-XJLJ202307) to F.Z., and a start-up grant from the University of Hong Kong to B.B.

## Author contributions

R.Z., F.Z., and B.B. conceived and designed the experiment. R.Z., G.J., and F.Z. selected the videos of studies 1 and 2. R.Z., X.G., L.W., and X.L. collected the data. R.Z. preprocessed the data for studies 1 and 2. R.Z., F.Z., and B.B. analyzed the data and were responsible for the interpretation of the data. X.G., T.X., F.Y., and X.S. provided important suggestions during the formal analysis. R.Z. drafted the manuscript. X.G. and T.X. provided suggestions during manuscript writing. F.Z. and B.B. revised the paper. F.Z. and B.B. contributed equally.

## Competing interests

The authors declare no competing interests.

## Notes

### Competing Interest Statement

The authors have declared no competing interest.

### Summary of Updates

We included several new analyses, six new datasets, and extended discussions and clarifications.

